# Buoyancy-driven sorting of synthetic cells for nanopore activity

**DOI:** 10.64898/2026.02.03.702772

**Authors:** Maja Lehr, Mattea Unger, Tobias Abele, Stefan J. Maurer, Dirk Flemming, Kerstin Göpfrich

## Abstract

The development of sorting strategies that directly report on functional activity remains a bottleneck in synthetic cell research. Current methodologies typically rely on sequential label-dependent probing, which limits throughput. Here, we introduce a label-free, buoyancy-driven selection strategy in which the mode of separation and the mode of decision-making are intrinsically linked, coupling pore activity directly to the synthetic vesicles’ internal density in a one-pot assay. In this system, sorting emerges intrinsically: Giant unilamellar vesicles (GUVs) that contain a dense medium sediment by default, while only those with functional transmembrane pores undergo solute exchange, leading to density equilibration and flotation. We exploit this principle to separate pore-active from non-functional GUVs without external markers or imaging-based readouts. Using protein pores and DNA origami and DNA tile nanopores, we demonstrate that buoyancy-driven separation enables parallel functional assessment of heterogeneous populations and supports flow-based enrichment of highly active synthetic cells. By directly linking molecular transport performance to GUV buoyancy, this approach collapses decision-making into the physical separation process itself, providing a scalable platform for screening, sorting, and evolving membrane pores in synthetic cell systems.

## 1 Introduction

Bottom-up synthetic biologists have become increasingly interested in sorting strategies for the directed evolution of synthetic cells – a paradigm shift from pure rational engineering to an approach that integrates evolution, as a fundamental principle of life, into the design process of synthetic cells^1 2^. While cell biology encompasses an arsenal of established sorting methodologies, bottom-up synthetic biology requires specific adaptations for its distinct chassis: the giant unilamellar vesicle (GUV). GUVs serve as widely used cell mimics to study membrane-associated functions, including nanopore-mediated transmembrane transport^3^.

Cell sorting and separation techniques can be organised along two largely orthogonal axes: the mode of decision-making and the mode of separation. While GUVs are compatible with many established methodologies, existing workflows typically decouple these two axes, necessitating sequential steps that limit overall throughput.

The mode of separation generally falls into two categories: displacement or trapping. Displacement-based methods rely on external forces to move cells or GUVs, most notably via electric fields in fluorescence-activated cell sorting (FACS)^4 5^ or flow-based separation in microfluidic channels^6^. Other modalities include optical tweezers^7^ or acoustic radiation forces^8^. Conversely, stationary approaches rely on immobilization – either through capture by molecular recognition (e.g., aptamer- or antibody-based capture of extracellular vesicles^9^, transient ligand interactions^10^) or spatial confinement by photopolymerization^11 12 13^, microwell arrays^14^ or photocleavable linkers^15^. While these methods are excellent for imaging, they often require complex “capture-and-release” mechanisms to transition from characterization to downstream recovery.

The primary bottleneck, however, lies in the mode of decision-making. In most synthetic cell platforms, sorting decisions are based on extrinsic features, such as fluorescent labels or visual morphological traits. This requires a serial interrogation of each candidate, where a sensor must first “decide” the vesicle’s fate before an actuator executes the separation. For populations with a very low hit rate, manual picking or site-specific photopolymerization^11^ can be adapted to individual GUVs, but these remain prohibitively low-throughput.

Transmembrane pore insertion is a key example for a desirable function in synthetic cellar systems as it enables communication, nutrient supply and waste removal^16 17 18 19^. Yet assessing the functionality of synthetic transmembrane pores is often limited to low throughput assays.

Currently, functional transmembrane pore insertion is typically monitored via fluorescent reporter transport, such as dye leakage assays^20^, dye influx assays enabled by confocal fluorescence microscopy^21^, single-channel ionic current recordings^22^, and analysing fixed states through electron microscopy^23^. While these techniques confirm pore insertion and activity, they predominantly probe individual vesicles or small subsets of a population, rely on potentially perturbative labels^24^, and provide limited access to population-level heterogeneity or insertion efficiency. As a result, functional screening and sorting of heterogeneous GUV populations remains challenging, particularly when pore insertion efficiencies are low.

To date, physical separation mechanisms for GUVs have been explored mainly for pu-rification^6^ rather than functional selection. There is a distinct lack of strategies where the decision-making is intrinsic to the separation itself. By coupling a biological function such as transmembrane transport directly to a physical property like buoyancy, the environment can perform the “computation” of fitness, allowing for a truly parallel, label-free sorting process.

Here, we propose a method for intrinsic self-sorting, where the sorting outcome emerges passively from the physical and functional properties of individual vesicles. By coupling nanopore activity directly to buoyancy-driven density equilibration, we bypass sequential probing and enable the label-free, one-pot selection of functional synthetic cells from heterogeneous populations.

The approach exploits a simple physical principle: when GUVs encapsulating a dense internal solution are placed in a lighter external medium, they sediment unless functional transmembrane pores permit solute exchange (Fig. 1). Pore-mediated transport passively equilibrates the internal and external densities, rendering vesicles neutrally buoyant and causing the full population to redistribute along the gravitational axis according to pore activity. Crucially, no active external decision process is required, as the orthogonal axes of separation mode and decision-making are intrinsically linked. This enables a comprehensive assessment of pore insertion efficiency and transport activity at the whole population level. We validate this concept using the well-characterised protein pore *α*-hemolysin, before applying the assay to DNA origami^25 18^ and DNA tile^26^ nanopores, which are known to produce highly heterogeneous GUV populations. By combining buoyancy-driven autonomous sorting with trapping of sedimented vesicles and flow-based collection of the pore-active GUVs, we establish a scalable, imaging-free functional readout and separation strategy for synthetic cells.

**Fig. 1.**
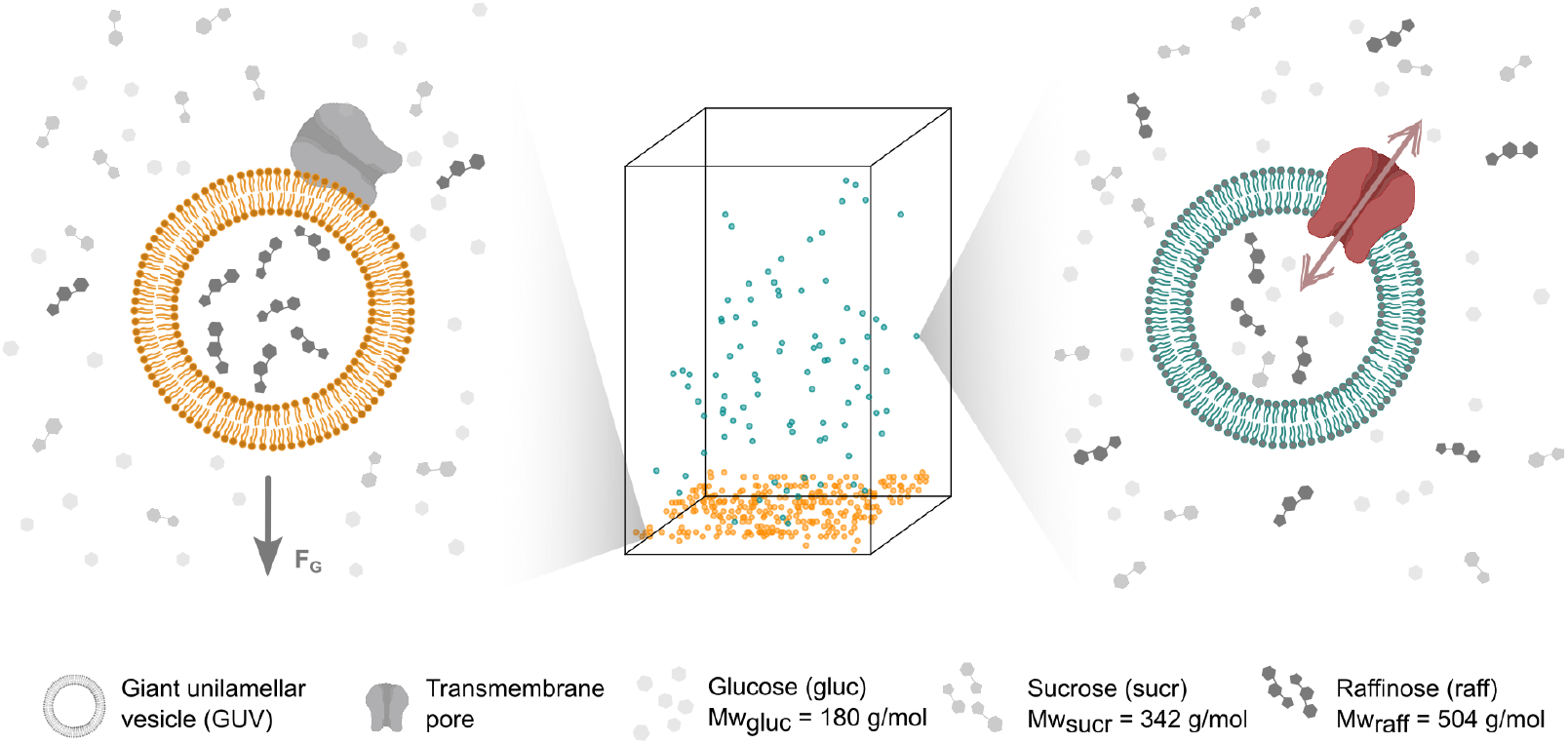
Concept of buoyancy-driven sorting of GUVs for nanopore activity. GUVs are first resuspended in a medium with a lower density. If a GUV cannot exchange the heavy internal solution with the external medium via transmembrane pores, the GUV sediments (orange, left). Functionally inserted transmembrane pores, on the other hand, allow the GUV to equilibrate the internal solution with the external medium, resulting in flotation (cyan, right).

## Main

### Probing sedimentation of GUVs

In order to implement a non-supervised buoyancy-driven sorting mechanism, we first need a detailed quantitative understanding of the expected timescales under conditions that are compatible with standard GUV experiments. GUVs are often produced in sugar-containing buffers. Thus, we chose sugars of different molecular weights, namely raffinose (Mw_raff_ =504 g*/*mol), sucrose (Mw_sucr_ =342 g*/*mol) and glucose (Mw_gluc_ =180 g*/*mol). We produced GUVs in raffinose as the heaviest of the sugars (Supplementary Fig. 1; Methods). When diluted in the same medium, the resulting GUVs exhibit minimal sedimentation (Fig. 2 **a**). Increased sedimentation is observed when the GUVs are diluted in a slightly lighter sugar solution (sucrose; Fig. 2 **b**), and the highest degree of sedimentation in the lightest sugar environment (glucose; Fig. 2 **c**). This is consistent with buoyancy expectations. Timelapse confocal microscopy confirms the progressive accumulation of GUVs at the bottom of a chamber equipped with microwells under these conditions within 30 min (Fig. 2 **d**). Similar behavior is observed when other high-molecular-weight sugars, such as maltotriose, were used for GUV formation (Supplementary Fig. 2).

**Fig. 2.**
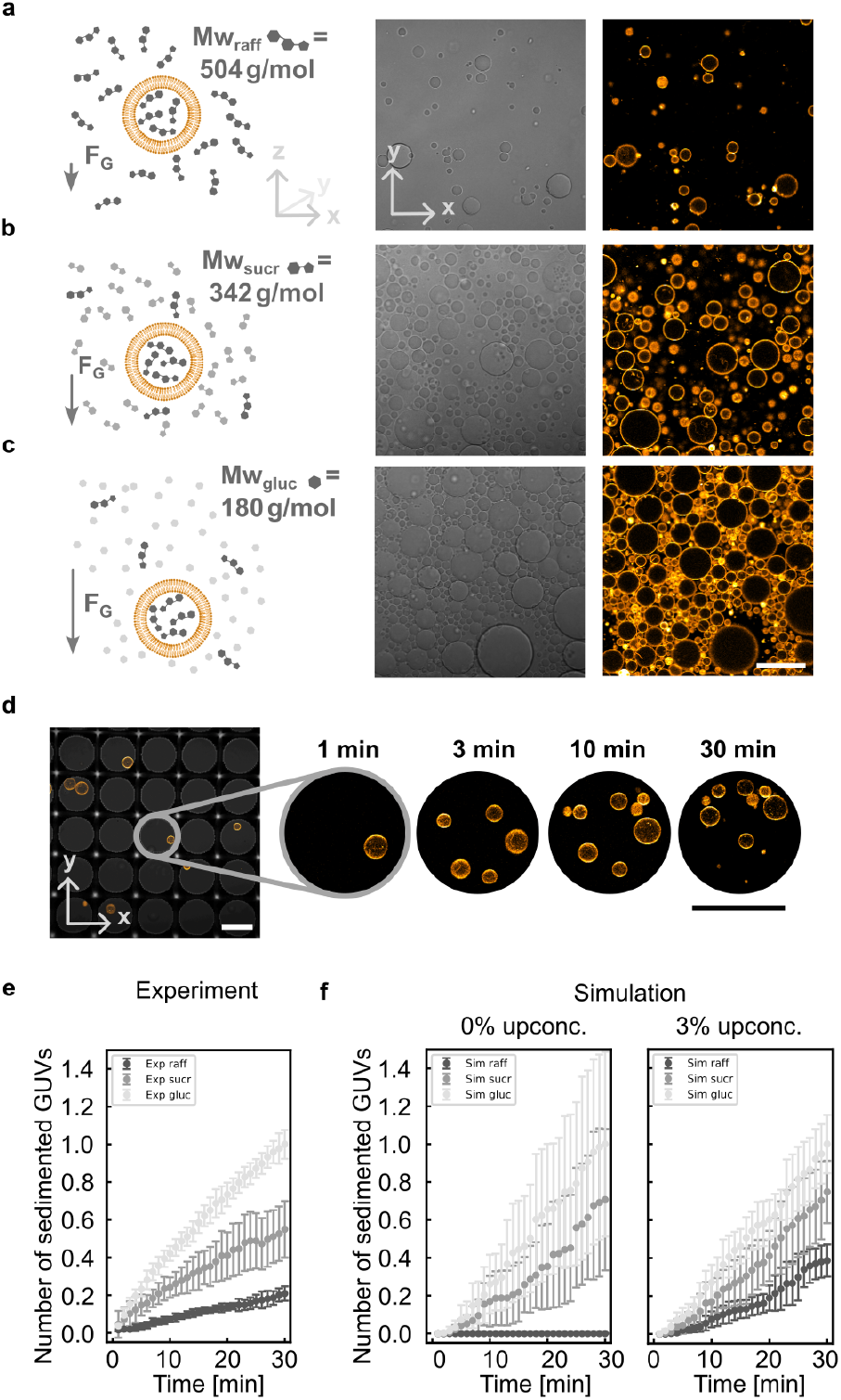
Buoyancy-driven sedimentation of GUVs. Immersion of raffinose-filled GUVs in different environments results in distinct sedimentation behavior. **a-c** Schematic representation (left) of GUV sedimentation, brightfield (middle, x-y-plane) and confocal image (right, x-y-plane, 99 % DOPC, 1 % Atto550-DOPE, *λ*_*ex*_ = 561 nm) close to the bottom of the observation chamber. Scale bar: 100 µm. Raffinose-filled GUVs mostly float in a medium containing purely raffinose (a), but they sediment if diluted in media containing components with a lower molecular weight, i.e. sucrose (b), or glucose (c). **d** Confocal images of sedimented GUVs in microwells. The zoom shows different time points in the sedimentation process as indicated. Scale bars: 100 µm. **e** Cumulative number of sedimented GUVs plotted over time in different sugars (*A*_img_ = 0.4 mm^2^, mean (circles) and standard deviation are shown, *n* = 3). **f** Simulation of the cumulative number of sedimented GUVs plotted over time in different sugars without any upconcentration of raffinose inside GUVs (left) and with an upconcentration of 3 % (right; mean (circles) and standard deviation are shown, *n* = 3).

Sedimentation of GUVs in lighter sugar solutions is widely employed as a washing strategy to remove lipid debris from the GUV solution^17^. In our system, raffinose-filled GUVs can be washed in sucrose and subsequently the sedimentation assay can be conducted in a glucose environment to increase the density difference. We identified overnight incubation as the optimal washing duration: longer times resulted in the accumulation of small lipid debris, while shorter times still showed an increase in GUV counts afterwards (Supplementary Fig. 3). Alternatively, centrifugation at 100 g for 60 min provided an effective washing step (Supplementary Fig. 3).

The number of sedimented GUVs was tracked over time by confocal microscopy (Fig. 2 **e**). Interestingly, we observe slow sedimentation of the raffinose-filled GUVs in raffinose, which cannot be explained by gravitational effects on the lipids alone but only by an upconcentration of the sugar during the electroformation process. The simulations clearly show no sedimentation of raffinose-filled GUVs in raffinose without upconcentration(Fig. 2 **f**, left).

By taking an upconcentration of raffinose into account, the simulation can be fitted to the experimental data, providing an estimate for an upconcentration of around 3 % compared to the surrounding medium during formation (Fig. 2 **f**, right; Supplementary Fig. 4, Supplementary Note 1). An accumulation of biomass is an important concept in the origins of life and is assumed to be a key driving factor during encapsulation and vesicle formation processes^27^.

### Buoyancy-driven sorting of GUV populations

Next, we performed proof-of-principle experiments to test whether a sedimentation assay can discriminate between permeabilised and non-permeabilised GUVs and, in principle, allow for sorting of these two populations. To characterise the sedimentation and buoyancy behavior of raffinose-filled GUVs in glucose with defined membrane permeabilities, we prepared two GUV populations labeled with distinct membrane dyes (Fig. 3 **a**). One population was incubated with *α*-hemolysin (cyan, *λ*_ex_ = 488 nm), a commonly used bacterial transmembrane pore that enables efficient transport of small molecules across the lipid bilayer. *α*-hemolysin has been used for nutrient supply and waste removal in synthetic cells, e.g. to initiate transcription via externally supplied triggers^17^ or to compare against synthetic DNA pores^20^. The second population remained pore-free to serve as a non-permeabilised control (orange, *λ*_ex_ = 561 nm). Following an incubation period to allow for pore insertion, the two populations were mixed and introduced into a 400 µm high observation chamber. Z-stack confocal imaging was performed using a long working distance objective, acquiring optical sections every 5 µm to minimise acquisition time while retaining sensitivity to small GUVs. The measured z-position of unpermeabilised GUVs (orange, *λ*_ex_ = 561 nm) reveals sedimentation to the bottom of the chamber of the entire GUV population (orange, *λ*_ex_ = 561 nm), while the z-positions of permeabilised GUVs of the other population remain scattered over the entire height of the chamber (cyan, *λ*_ex_ = 488 nm; Fig. 3 **b, c**; Supplementary Movies 1-4). Confocal imaging at defined z-positions reveals that GUVs located near the bottom of the chamber do not show dye influx, whereas GUVs capable of exchanging solutes with the surrounding medium due to the presence of pores display clear influx of the fluorescent probe (purple, *λ*_ex_ = 640 nm; Fig. 3 **b**). GUVs that do exhibit dye influx are also capable of releasing raffinose, leading to flotation. While unpermeabilised GUVs accumulate at the bottom of the observation chamber, permeabilised GUVs are distributed across all z-planes.

**Fig. 3.**
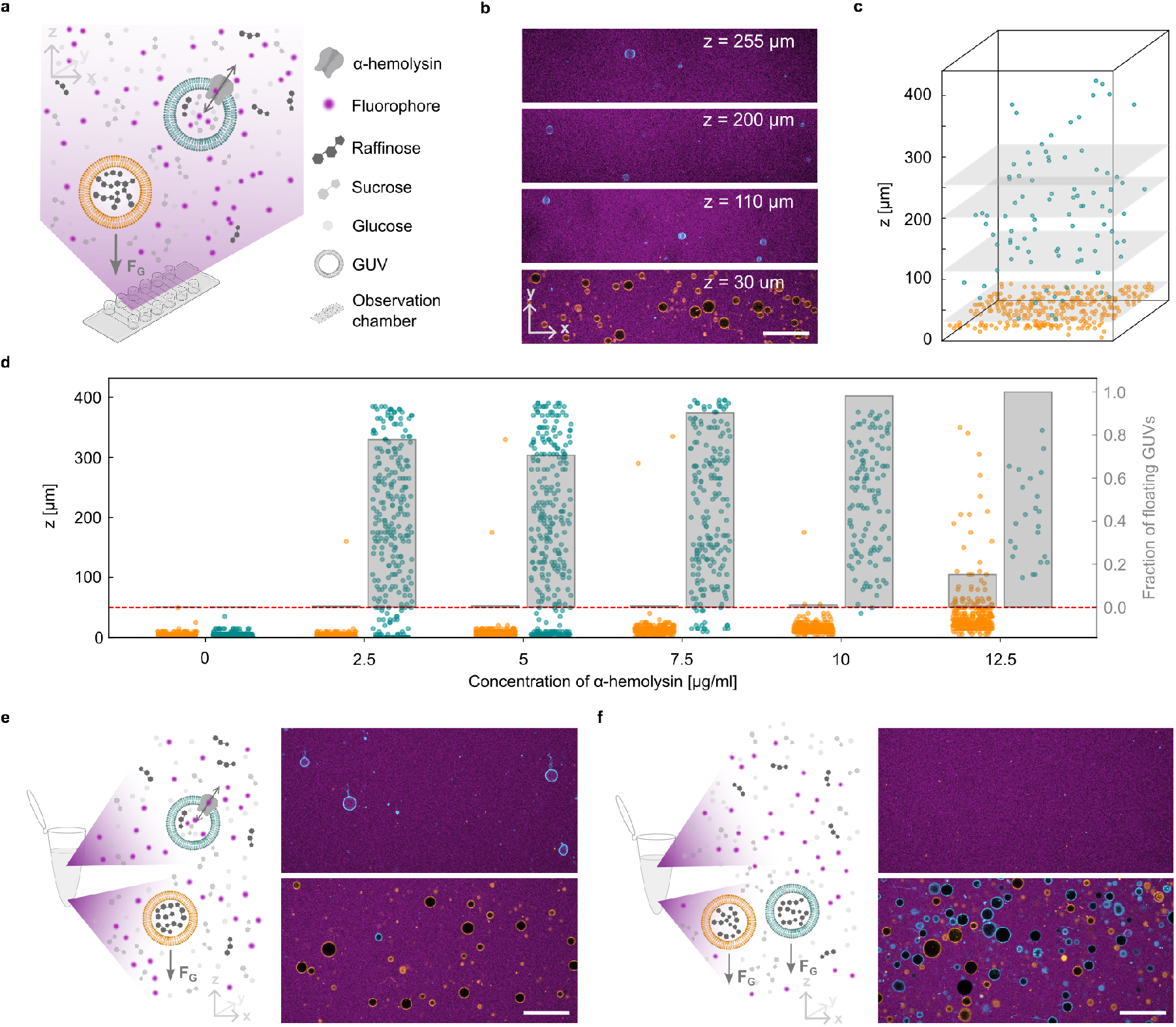
Proof-of-principle sedimentation assay of GUV populations with and without membrane pores. **a** Schematic representation of two populations of GUVs without *α*-hemolysin (orange) and with *α*-hemolysin (cyan) in a 400 µm high observation chamber. **b** Confocal images of GUVs in the upper layers (cyan, 99 % DOPC, 1 % Atto550-DOPE, *λ*_ex_ = 561 nm, 0.625 µg*/*ml *α*-hemolysin) and lowest layer (orange, 99 % DOPC, 1 % Atto488-DOPE, *λ*_ex_ = 488 nm) immersed in influx dye (purple, Alexa Fluor 647, *λ*_ex_ = 640 nm) at indicated z-positions (from top to bottom *z* = 255 µm, *z* = 200 µm, *z* = 30 µm). Scale bar: 100 µm. **c** Z-position of pore-free GUVs (orange data points) and GUVs with 0.625 µg*/*ml *α*-hemolysin (cyan data points) mixed in one chamber, represented over a schematic x-y-scale. **d** Z-position of pore-free GUVs (orange data points) and GUVs with increasing *α*-hemolysin concentration (cyan data points) and fraction of floating GUVs (gray bars). **e,f** Schematic representation of sedimentation assay in a test tube with mixed populations of GUVs with and without pores. Respective confocal images show solution taken from the top part (top) and bottom part (bottom) of the test tube after 3 h incubation at 37 °C, the cyan GUV population contains 5 µg*/*ml *α*-hemolysin (e) and control (f). Scale bars: 100 μm.

To determine the effect of *α*-hemolysin on the fraction of floating GUVs, we tested concentrations from 0 to 12.5 µg*/*ml. The maximum of fraction of intact floating GUVs (98 %) was observed at 10 µg*/*ml of *α*-hemolysin, indicating efficient pore insertion. However, at 12.5 µg*/*ml, the *α*-hemolysin–treated GUVs showed signs of membrane damage, including rupture and a marked decrease in the number of intact vesicles. At this concentration, the untreated population also displayed substantial dye influx and flotation, likely caused by free pores in solution (Fig. 3 **d**). Acquisition time was minimised at optimised field of view to buffer the variance of the sample (Supplementary Fig. 5).

We also prepared a 1:7 and a 7:1 mix in addition to the 1:1 mix of permeabilised to unperme-abilised GUVs, confirming efficient separation along the gravitational axis and high sensitivity of the assay independent of the fraction of pore-active GUVs (Supplementary Fig. 6).

To test whether the sorting mechanism can be scaled up to large populations, we assessed the flotation and sedimentation properties of the two GUV populations using an alternative assay where sedimentation was performed in 1.5 ml reaction tubes. After separate incubation, subsequent mixing and equilibration, GUVs were collected from the top and bottom of the tube, respectively. As during the microwell-based sorting experiments, permeabilised GUVs predominantly occupy the top fraction of the reaction tube, while non-permeabilised GUVs sediment to the bottom (Fig. 3 **e**). When the second population (cyan, *λ*_ex_ = 488 nm) was not treated with *α*-hemolysin, both fluorescently labeled populations were found exclusively at the bottom of the 1.5 ml tube (Fig. 3 **f**, Supplementary Fig. 7). This demonstrates that our assay is in principle capable of sorting millions of GUVs in a simple one-pot step and further scale-up for sedimentation of large volumes should be possible – compatible with large-scale directed evolution experiments.

### Screening for DNA pore activity

Next, we tested whether buoyancy-driven separation can resolve the functional heterogeneity of reconstituted DNA origami pores, which have variable assembly yields and insertion efficiencies^28 19 18^.

Despite ongoing efforts to improve insertion efficiency^29^, synthetic DNA pores do not insert into all GUVs, leading to heterogeneous GUV populations. It would be desirable to compare the fraction of functional insertions for different pore designs and to sort functional from non-functional GUVs for downstream experimentation. We used a DNA origami pore with a 12 helix bundle (12HB) architecture which has been designed and characterised earlier (Fig. 4 **a**)^25 30 18^. It has a length of 63 nm, an inner pore diameter of 7.3 nm, an outer diameter of approximately 12 nm and carries 24 positions for modification with hydrophobic membrane anchors. We confirm the correct assembly of the 12HB DNA origami pore with negative stain transmission electron microscopy (TEM), super-resolution microscopy and gel electrophoresis (Fig. 4 **b,c**; Supplementary Figure 8, 9 and 10). Ensemble averaging clearly reveals the correctly assembled stem of the pore as well as a smeared density on the left side of the stem, corresponding to the flexible fluorophore-bearing scaffold loop (Fig. 4 **c**). Additionally, the blurred ends of the pore stem suggest that the ends of the 12HB are fraying. Confocal imaging confirms the efficient attachment of the cholesterol-modified 12HB DNA origami pore to the GUV membrane (Fig. 4 **d**), but note that attachment does not necessarily correspond to functional insertion.

**Fig. 4.**
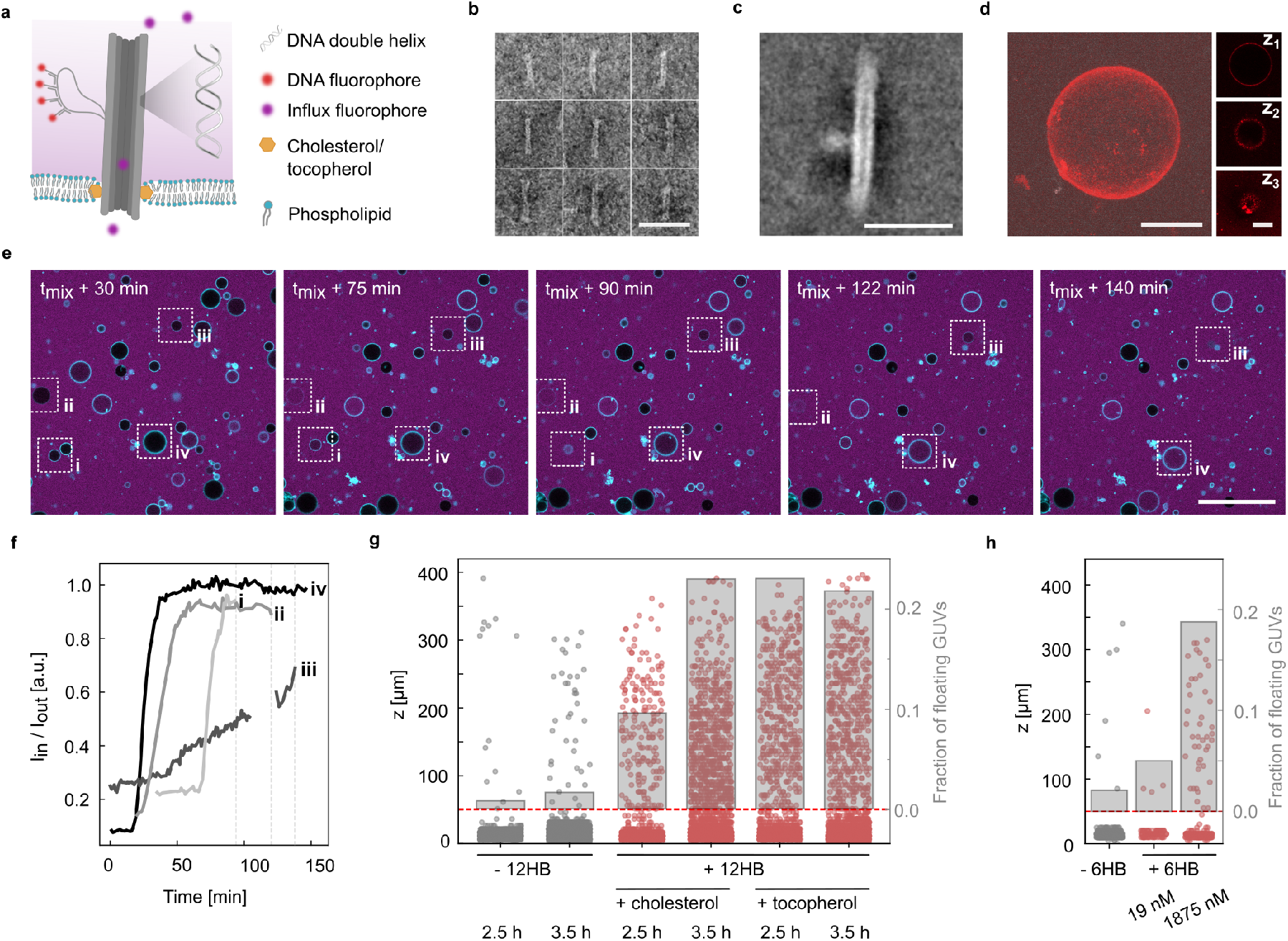
Sedimentation assay to assess the efficiency of a 12HB DNA origami pore. **a** Schematic representation of the 12HB DNA origami pore with its membrane anchors. **b** Negative stain transmission electron microscopy (TEM) of the 12HB DNA origami pores. Scale bar: 100 nm. **c** Ensemble averaging of the TEM micrographs. The lower density next to the stem corresponds to the scaffold loop bearing the fluorophores. Scale bar: 50 nm. **d** Confocal z-projection of a GUV covered by Atto647-labeled 12HB DNA origami pores and confocal planes at different z-positions (*z*_1_, *z*_2_, *z*_3_, *λ*_*ex*_ =640 nm). Scale bars: 10 µm. **e** Confocal timelapse images of GUVs (cyan, 99 % DOPC, 1 % Atto488-DOPE, *λ*_*ex*_ =488 nm) equipped with 12HB DNA origami pores. GUVs initially sit at the bottom observation chamber. DNA origami pore insertion causes dye influx (purple, Alexa Fluor 546, *λ*_*ex*_ =546 nm) followed by floating, hence perforated GUVs disappear from the imaged confocal plane (white boxes). Scale bar: 100 µm. **f** Dye influx (internal over external intensity) over time for GUVs that show 12HB DNA origami pore insertion (**i** to **iv** as indicated in (e)). **g** Z-position of GUVs incubated without and with 20 nM 12HB DNA origami pores functionalised with 24 cholesterol or 24 tocopherol membrane anchors and at different incubation times. The fraction of floating GUVs is indicated in grey, revealing that up to 23 % of GUVs show pore insertion. **h** Z-position of GUVs incubated without and with 20 nM and 2 µM of the 6HB DNA tile pore functionalised with three cholesterol tags.

Next, we used the sedimentation assay to quantify the insertion efficiency of the 12HB DNA origami pore into the GUVs. GUVs were first washed by sedimenting them in sucrose and subsequently immersed in a glucose environment. Note that we observe that raffinose-filled GUVs are more prone to burst at a PVA-coated glass surface of an observation chamber in presence of magnesium ions. PVA-coating on silanised glass slides efficiently prevents GUV disruption (Supplementary Fig. 11, see Methods).

Upon addition of DNA origami pores, some GUVs exhibited dye influx, then they begin to float and finally disappear from the confocal plane. This onset of flotation of the GUVs occurs without signs of membrane budding or tubulation, suggesting that the observed behavior is due to buoyant flotation rather than vesicle deflation or ongoing lysis (**i-iv**, Fig. 4 **e,f**; Supplementary Movie 5).

In samples incubated with the 12HB DNA origami pores, we anticipate a heterogeneous population comprising GUVs that remain at the bottom and those that float in z. Consistent with this expectation, the fraction of floating GUVs is below 2 % in control samples lacking the 12HB pore, but increases to over 20 % in samples incubated for at least 3.5 h with 20 nM of the 12HB DNA origami pore and cholesterol anchors (Fig. 4 **g**). An increase in the fraction of floating GUVs can be observed over the course of up to 7 h (Supplementary Fig. 12 **a,b**), but it clearly remains below the fraction of permeabilised GUVs observed in the presence of 22 nM *α*-hemolysin (5 µg*/*ml) pores. When using tocopherol as membrane anchor instead of cholesterol, the same fraction of floating GUVs is reached after only 2.5 h of incubation, indicating faster pore incorporation. This could potentially be explained by a lower propensity for aggregation of the tocopherol-tagged pores. The pre-incubation of the cholesterol DNA strand with GUVs at 35 °C for 10 min does not infer with the GUV integrity, but shows no benefit for the fraction of floating GUVs either (Supplementary Fig. 12 **c**). Raising the temperature to 30 °C increases the fraction of floating GUVs, whereas cooling to 4 °C reduces it remarkably (Supplementary Fig. 12 **d**). These observations offer guidance for the choice of parameters when working with DNA origami pores. Mildly elevated temperatures and incubation times of several hours increase insertion efficiency.

Note that while magnesium ions themselves can destabilise membranes and lead to flotation, this effect disappears when cholesterol-tagged DNA is added (Supplementary Fig. 13).

Next, we compared insertion efficiencies of different DNA pore architectures. While the 12HB pore uses the scaffolded DNA origami technique, another popular class of DNA pores are made from DNA tiles^31 22 32^. They are annealed without a scaffold from 2-12 individual oligonucleotides and most commonly adopt a 6-helix bundle (6HB) architecture. Here, we use a 6HB tile pore assembled from six strands, three of which carry a terminal cholesterol modification^26^. It has a length of 9 nm, an outer pore diameter of approximately 5 nm and an inner pore diameter of approximately 2 nm compared to 63 nmx12 nmx7 nm (length, outer and inner diameter) for the 12HB origami pore.

Because of the scaffold-free design, DNA tile pores can readily be annealed at micromolar concentrations without the need for subsequent purification. We thus performed the sedimentation assay with the 6HB pore at the same concentration as perviously with the 12HB pore (20 nM) as well as at a 100× higher concentration of 2 µM. At 2 µM, we obtain almost 20 % of floating GUVs – similar to the 12HB pore – while there is only a negligible fraction of floating GUVs for the lower concentration (Fig. 4 **h**, Supplementary Fig. 14). This is not surprising when considering the free energy balance for insertion of the two pore variants: the 6HB pore carries an eightfold lower number of cholesterol anchors compared to the 12HB pore, which cannot be compensated by the smaller pore diameter. From previous coarse-grained molecular dynamics simulations^33^ we can deduce that the free energy for insertion of the 6HB with three cholesterol anchors is close to net 0 compared to −1000 kJ*/*mol for 12HB containing 24 membrane anchors. Nevertheless, minimal pore constructs are capable of inducing buoyancy-driven separation at high enough concentrations (Fig. 4 **h**).

### Harvesting of sorted pore-active GUVs

So far, we have observed that GUVs containing natural protein pores as well as synthetic DNA pores float, while non-functional GUVs settle. Next, we set out to make use of this principle to implement a high-throughput automated sorting mechanism that allows to collect pore-active GUVs efficiently – without any optics, fluorescent tagging or the need for complex active sorting processes. Our setup consists of a microwell chamber connected to a peristaltic pump (Fig. 5 **a**). Microwells are well established to immobilise single cells on a chip^34 35^, but are also widely used as traps for GUVs^13 14^.Our microwell chamber was prepared in-house by photopolymerization with structured illumination from a digital mirror device (for details see Methods and Supplementary Fig. 15-17). As a bench-top fluidic setup, it remains easy to use and allows also flexible manual pipetting.

**Fig. 5.**
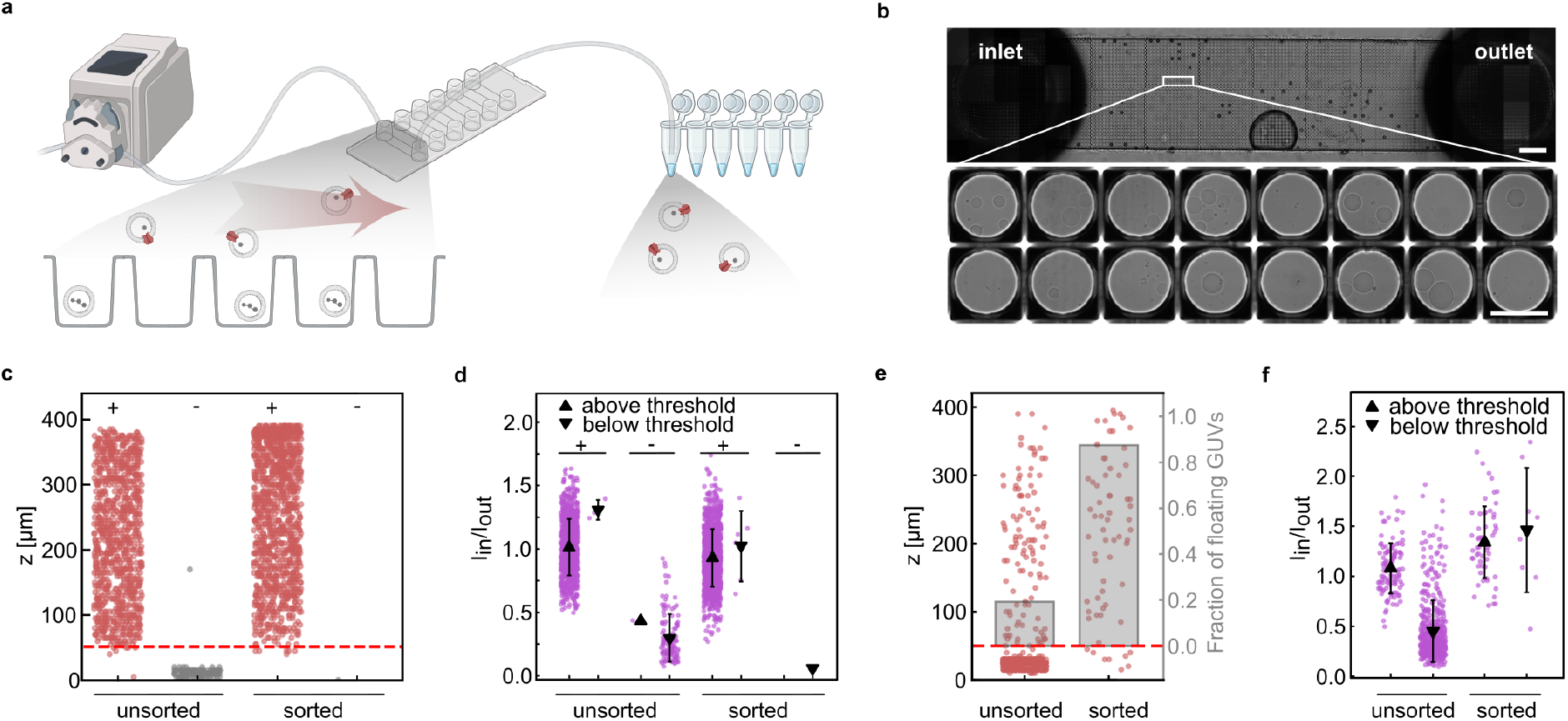
Buoyancy-driven sorting of pore-active GUVs. **a** Schematic representation of the automated sorting setup. A peristaltic pump flushes out pore-active GUVs that are subsequently collected in a reaction tube. GUVs without pores settle in microwells inside the channel. **b** Brightfield microscopy image of microwells at the bottom of the chamber immobilise GUVs without pores. Scale bars: 1 mm (top), 100 µm (bottom). **c** Measured z-position of a mix of prepared populations of GUVs incubated with and without *α*-hemolysin (+, -) before flushing them through the microwell chamber (unsorted) and after sorting (sorted). Up to 3864 GUVs were reimaged after collection. **d** Mean and standard deviation of inner over outer intensity of influx dye for data shown in (c) above and below z-threshold (upward and downward pointing triangle) for GUV populations incubated with and without *α*-hemolysin (+, -). The *α*-hemolysin-containing GUVs show dye influx (99.9 % before and 98 % after sorting, defined as *I*_*in*_ *>* 0.5*I*_*out*_). Only 17 % of the GUVs without *α*-hemolysin show dye influx before sorting, none is observed after sorting. **e** Measured z-position of GUVs incubated with 12HB DNA origami pore before flushing them through the microwell chamber (unsorted, 19 % floating) and reimaged collection after sorting (sorted, 88 % floating). Up to 670 GUVs were reimaged after sorting. **f** Mean and standard deviation of inner over outer intensity of influx dye for data shown in (e) above and below z-threshold (upward and downward pointing triangle) for GUV populations incubated with and without *α*-hemolysin (+, -). Before sorting 41 % of GUVs show dye influx (defined as *I*_*in*_ *>* 0.5*I*_*out*_), after sorting 98 % show dye influx.

GUVs were first introduced into the chamber by pipetting such that the GUVs do not need to be circulated through the pump head which could cause mechanical disruption. Afterwards, an applied air pressure by the peristaltic pump moves the floating fraction out of the chamber for collection without dilution of the sample.

Figure 5 **b** shows the microwell architecture visualised by brightfield microscopy, but note that microscopy is not required for successful sorting. Height and width of the microwells were chosen based on flow simulations^36 37 38^ such that sedimented GUVs experience stable immobilization at the bottom of the wells (Supplementary Movie 6), while GUVs located only slightly higher are efficiently entrained by the applied flow in the upper region of the microwell (Supplementary Fig. 17, Supplementary Movie 7). As a result, even small differences in vertical position translate into a robust binary outcome—either immobilization or export by flow. Non-permeabilised GUVs remain trapped within the wells, whereas pore-active GUVs are selectively flushed out and collected downstream in a reaction tube, yielding a highly precise and pure pore-active population.

The harvested GUVs can be subsequently re-analysed by transferring them back to an imaging chamber, either manually or by directly connecting the collection tubing to an imaging setup (Supplementary Fig. 18).

With the current microwell chamber design, consisting of approximately 6x 5000 wells, we can immobilise approximately 6x 25,000 GUVs. To benchmark the sorting process, we prepared two populations of GUVs with and without *α*-hemolysin and mixed them in a 1:1 ratio, hence we start out with a ratio of 50 % pore-active GUVs. After sorting, the fraction of pore-active GUVs is 99.2 % (Fig. 5 **c**, Supplementary Fig. 19; 0.09 % false positive). We show that 98 % of sorted GUVs show influx (Fig. 5 **d**) confirming the functionality of the pores.

We also tested whether the manufactured heterogeneity in the population mix of 1:7 and 7:1 can be sorted. We observe successful sorting, and the recovery of the expected fractions (Supplementary Fig. 20).

For DNA origami pores, this approach enables the enrichment of a heterogeneous population initially containing less than 20 % floating GUVs to over 98 % functional GUVs showing dye influx (Fig. 5 **e,f**), effectively isolating the most potent candidates with well-inserted DNA origami pores (Supplementary Fig. 18). This demonstrates that the assay’s high sensitivity to buoyancy-driven separation of a full GUV population along the gravitational axis is effectively converted into high sorting precision for functional nanopore insertion.

## Conclusion

We present a buoyancy-driven separation strategy that enables functional screening and sorting of heterogeneous GUV populations based on transmembrane pore activity. By coupling solute exchange through membrane pores to density equilibration, GUVs autonomously redistribute along the gravitational axis, allowing pore-active GUVs to be distinguished from non-functional ones without labels, reporters, or imaging-based readouts. No external, sequential decision-making is required, as the physical properties of individual vesicles intrinsically determine their sorting behavior, directly linking pore activity to spatial separation. We demonstrate that the assay discriminates between pore-active and non-permeable GUVs by reconstituting protein pores as well as DNA origami and DNA tile pores. The resulting flotation of pore-active GUVs provides a direct, population-level functional readout and enables parallel assessment of pore performance within mixed populations, supporting the enrichment of functional pore variants through flow-based separation. Importantly, the assay operates under mild conditions and is compatible with diverse pore architectures, buffer compositions, and incubation protocols.

Beyond characterization, this platform establishes a foundation for selection-based workflows in synthetic cell engineering. Libraries of pore variants could be screened in a one-pot format, followed by enrichment, recovery, and downstream molecular analysis. With our current microwell format we have already demonstrated that we can sort 150, 000 GUVs at once, a one-pot format would increase that number to 10^6^ GUVs – which is often sufficient for directed evolution approaches. Scale-up to batch sorting of large volumes should be feasible.

Importantly, this work highlights the broader relevance of buoyancy effects, extending beyond pore performance screening. In particular, the platform is applicable to synthetic cell engineering scenarios that rely on the external supply of reagents across membranes, enabling quantitative readouts of reaction efficiency. For example, RNA transcription within vesicles has been shown to be initiated by the transmembrane supply of magnesium ions and nucleotides^17^, such that candidates with higher membrane perforation rates exhibit increased RNA production.

The findings presented here are also highly relevant for dye influx assays that are routinely used in the nanopore community. In these assays, it is common practice to sediment GUVs in a density contrast and perform timelapse imaging of the confocal plane close to the bottom of the observation chamber^18 19 21^. Given our findings, it is conceivable that these assays underestimate pore activity, because pore-active GUVs disappear from the imaged confocal plane due to solute exchange. Such effects may remain unnoticed and are important to consider for future experiments.

Note that the passive redistribution driven by buoyancy rather than active self-organization or collective interactions, is a concept that resonates with scenarios proposed for the origins of life, where heat flows and related gradients are thought to have driven the accumulation of dilute biomolecular components into increasingly concentrated compartments^27^. In such settings, compartments that accumulate sufficient mass would sediment, yet this very accumulation could promote changes in membrane properties and the emergence of pores.

More broadly, buoyancy-driven separation offers a physically grounded, prebiotically plausible mechanism by which functional membrane permeability could confer a selective advantage. The “floating of the fittest” mechanism is directly linking transport activity to spatial separation from their original environment and passive transport into new environments where further development and selection can occur to ensure survival. By collapsing decision-making into the physical behavior of individual vesicles, this strategy exemplifies how intrinsic sorting mechanisms can be relevant to advance origins of life and synthetic cell research alike.

## Methods

### GUV formation

The electroformation of the GUVs was conducted as reported in Jahnke & Illig et al.^18^. 1,2-Dioleoyl-sn-glycero-3-phosphocholin (DOPC), Atto 488 1,2-Dioleoyl-sn-glycero-3-phosphoethanolamine (488-DOPE), Atto 550 1,2-Dioleoyl-sn-glycero-3-phosphoethanolamine (550-DOPE) or 1,2-dioleoyl-sn-glycero-3-phosphoethanolamine-N-(lissamine rhodamine B sulfonyl) (ammonium salt) (LissRhod PE) were purchased from ATTO-TEC or Avanti Research. The lipids were mixed in the following ratios, if not stated otherwise, 99 mol% DOPC and 1 mol% fluorescent lipids at a total concentration of 5 mM in chloroform. The mix was prepared and stored at *−*20 °C for several weeks. The conductive side of indium tin oxide (ITO) coated glass slides was identified with a multimeter. The lipid mix was gently spread onto the conductive side with a 20x20 mm glass cover slip by hand allowing homogeneous layering of the lipids. ITO-slides were purchased from Nanion Technologies GmbH. The lipid film was dried in a desiccator while the vacuum pump was on for 10 min. Then, the vacuum was kept for additional 30 min or longer before the slides were retrieved. Subsequently, a rubber ring was covered with silicon grease and then positioned at the most homogeneous looking area of the lipid spread. The ITO slide with the rubber ring was already mounted into the Vesicle Prep Pro device (Nanion Technologies GmbH) before the internal solution (280 mOsmol*/*kg raffinose or sucrose) was preheated for 10 min to 50 °C and then 270 µl were added to the rubber ring, and the second glass slide was added with the conducting side pointing towards the solution. The Vesicle Prep Pro internal standard protocol was run for 138 min at 3 V and 5 Hz. Afterwards, GUVs were transferred on top of 800 µl of washing solution (280 mOsmol*/*kg sucrose or glucose) in a 1.5 ml tube and they kept in the fridge overnight for sedimentation or centrifuged for 1 h at 100 g (Centrifuge 5810, S-4-104 Rotor, Eppendorf). Subsequently, 200 µl of washed GUV solution was collected carefully from the bottom of the tube and transferred to another storage tube, stored at 4 °C for up to 4 weeks and gently mixed before use.

### Silanisation of glass slides

For silanisation, 24 mmx60 mmx170 µm glass slides (No. 1.5H, borosilicate, Marienfeld) were sonicated for 15 min in MilliQ water and ethanol, dried under nitrogen, and plasma-activated for 35 s at 200 W in 0.4 mbar oxygen (300 SemiAuto, PVA TePla AG or ZEPTO, diener) in an aluminum glass slide rack. In a teflon holder, a maximum of 20 glass slides are immersed in an approx. 300 ml toluene bath with 2.5 vol% 3-(Trimethoxysilyl)propylmethacrylate (TMSPMA; Sigma-Aldrich), covered with a glass lid and sealed with teflon tape and aluminum foil. The next day, slides in the teflon holder were washed in ethanol and MilliQ water and slowly removed from the washing bath to obtain dry surfaces. Slides were then stored between kimtech wipes for up to 3 weeks.

### Confocal fluorescence microscopy

Confocal microscopy images were acquired using a confocal laser scanning microscope LSM 900 (Carl Zeiss AG), 20× (Plan-NEOFLUAR 20×/0.5 Air M27, Carl Zeiss AG) or 63× objective (Plan-Apochromat 63×/1.4 Oil DIC M27). Experiments were performed at room temperature unless stated otherwise. The pinhole aperture was set to one Airy Unit. To reduce acquisition time while maintaining enough field of view size to buffer the variance of the sample, imaging parameters were optimised by lowering the spatial resolution down to 512×512 pixels, thereby accelerating collection of four tiles of the 80-slice z-stacks per condition (Supplementary Fig. 5). Under these optimised conditions, imaging five experimental conditions in four channels (two membrane dyes, one influx dye, and bright-field) took approximately *≈*1.5 h. To average out time-dependent effects in the sample, imaging was performed in a bidirectional sequence: conditions 1–5 were imaged in forward order for half of the required tiles, followed by imaging conditions 5–1 in reverse order for the remaining tiles.

### Sedimentation assay

First, a 400 µm high observation chamber with 6 channels composed of a sticky-Slide VI 0.4 (ibidi) and either a silanised (in case of presence of magnesium ions or printed structures) or an unprocessed glass slide (otherwise; Supplementary Fig. 11) was prepared. After assembly, the chamber was coated with filtered (0.22 µm sterile PES membrane filter, TPP) 5 wt% PVA (Sigma Aldrich) for up to 5 min, if not stated otherwise. Washed GUVs were mixed 1:10 in settling solution with 50 µg*/*ml influx dye (Alexa Fluor 647 NHS Ester or Alexa Fluor 546 NHS Ester, Sigma Aldrich) and *α*-hemolysin, 12HB DNA origami pore, 6HB DNA tile pore, cholesterol DNA or other ingredients as stated. 40 µl GUV solution were added to the channel. The conditions were imaged from bottom to top, 5 µm distance between z slices, preferably consecutively from condition 1 to 5 and back to minimise any time-related bias between conditions.

### Image data analysis

Image analysis was performed in Python (v3.10.14) and standardised on 512×512 pixel images. Multichannel tiled z-stack.czi files were imported and converted into individual image slices. Contrast was enhanced per slice using contrast-limited adaptive histogram equalization (CLAHE). Candidate GUVs were segmented by adaptive thresholding, followed by morphological filtering to remove noise (MorphologyEx), distance transform–based detection of local maxima, and watershed segmentation. Segmented objects were filtered based on geometric criteria (size, eccentricity, and border contact). For each GUV, the slice with the largest apparent radius across the z-stack was defined as the vesicle center. Segmentation results were validated by overlaying detected vesicle contours onto the original image data, and slice-wise images were saved.

### DNA pore assembly

The DNA origami used in this work was previously introduced by Jahnke & Illig et al.^18^ as a membrane spanning—DNA Origami Signaling Unit— enabling the transport of small molecules. The design is based on a structure by Mohammed et al.^25^, originally used as a seed for DNA nanotube assembly^39^ and later for transmembrane transport^30^. The DNA origami consists of an M13mp18 scaffold (tilibit) and in total 220 staple strands assembled into a 12-helix bundle cylindrical structure (12HB). Assembly was performed in 50 or 100 µl volumes using 1× TAE buffer with 12 mM MgCl_2_. Reaction mixtures contained 16 nM p7249 scaffold (100 nM stock, tilibit), 160 nM premixed staple strands (1 µM in 1× TE), 160 nM fluorophore-attachment strands, 16 µM Atto647-labeled strands (Biomers, HPLC purified), and 320 nM of two adapter-strand mixes. Unmodified strands were purchased from IDT (standard desalting). Samples were heated to 90 °C for 10 min and cooled either to 45 °C or 55 °C at 1 °C*/*min, or to respective temperatures for condition screening. They were held at 45 °C for 1 h (previous protocol adapted from^25^) or at 55 °C for 10 h (optimised protocol), then cooled to 32 °C at 0.1 °C*/*min in a thermocycler (C1000 Touch Thermal Cycler, Bio-Rad). The protocol is specified for each experiment. 12HB samples were stored at 4 °C for 7 days. Before use, 12HB was purified by spin filtration (Amicon, 100 kDa). Filters were equilibrated with 500 µl folding buffer (1× TAE, 12 mM MgCl_2_) and centrifuged at 4 °C for 5 min at 11.700 g. Subsequently, 450 µL folding buffer and 50 µL folded 12HB were added and centrifuged under the same conditions, followed by two identical wash steps with 450 µL folding buffer. 12HB concentrations were determined by measuring absorbance at 260 nm (NanodropOne, Thermo Scientific). For functionalization with membrane anchors, two stock solutions of single stranded DNA (100 µM in 1xTE, 3-prime and 5-prime modified with cholesterol, Biomers, HPLC purified) were heated to 90 °C for 2 min and then incubated with 12HB at a 0.02:0.02:0.96 volume ratio for 10 min at 35 °C and 200 rpm.

The 6HB DNA pore was assembled by heating the six strands at an equimolar mixture of 3.4 µM to 95 °C for 5 min in 1xTAE with 500 mM KCl and optionally with 10 mM MgCl_2_ followed by cooling down to 4 °C at a rate of 0.25 °C per min (total volume 100 µl).

### Negative stain electron microscopy

Negative staining, data collection, and processing were performed as described previously^40^. In brief, 5 µl sample was applied to a freshly glow-discharged grid with a 6-8 nm thick layer of continuous carbon. After incubation for 60 s the sample was blotted on a filter paper and quickly washed with three drops of water. Samples on grids were stained with 3% aqueous uranyl acetate and dried. Images were acquired using a ThermoFisher Talos L120C electron microscope equipped with a Ceta 16 M camera, operated at 120 kV. The micrographs were acquired at a nominal magnification of 28,000x (calibrated pixel size of 5 Å) or 45,000x (calibrated pixel size of 3.3 Å) using EPU software (Thermo Fisher Scientific). For 2D classification 1800 particles were selected using the cryoSPARC blob picker^41^ and extracted with a box pixel size of 384 pixel. Image processing was carried out using the IMAGIC-4D package^42^. Particles were band-pass filtered, normalised in their gray value distribution, and mass centered. 2D alignment, classification, and iterative refinement of class averages were performed as previously described^43^.

### Agarose gel electrophoresis

The assembly of the 12HB was evaluated using agarose gel electrophoresis. Gels were prepared at 0.7% agarose in 1× TBE buffer containing 12 mM MgCl_2_, unless stated otherwise. The same buffer was used as both the casting and running buffer. For each lane, 10 µl of the 12HB sample was mixed with 6× tilibit loading dye and loaded alongside a Tridye 1 kb Plus DNA ladder (New England Biolabs, MA, USA) as a reference. Electrophoresis was performed at 60-70 V for 3–5 h. Following the run, gels were stained in a 3× GelRed (Sigma-Aldrich) solution for 20 min and subsequently imaged using a Bio-Rad ChemiDoc MP Imaging System.

### Microwell-based sorting

Sorting chambers were prepared by bonding a multichannel slide (sticky-Slide VI 0.4, ibidi) onto a silanised glass substrate. A photocurable resin (Bio-Med Clear, Liqcreate) was then introduced into the channels. Microwells were printed using a custom-built photoprinting setup comprising a Polygon digital micromirror device (DMD) pattern illuminator (Polygon 1000, MIGHTEX) coupled to an inverted fluorescence microscope (Axio Observer 7; Carl Zeiss AG). Binary illumination masks defining circular or square microwells (80 µm to 100 µm lateral dimension) were generated using custom scripts written in Python (v3.12.4). The printing process was controlled through a custom macro written in the ZEISS ZEN Macro Environment. The focus was manually set to the bottom of the glass slide at 12 positions for each channel and saved as support points, from which the microscope could interpolate the correct focus for any position within a channel. Patterning was performed using a 385 nm LED (MIGHTEX) connected to the DMD, with the 5× objective (Plan-Apochromat 5x/0.16 M27, Carl Zeiss AG), with each tile exposed for 200 ms. To cover the full channel area, adjacent 5× fields of view were sequentially exposed by automated tiling throughout the entire sample area. The microwell channel was washed in ethanol and MilliQ. Subsequently, 50 µL of 5 wt% PVA were added and the entire slide is desiccated to extract remaining air from the microwells. The applied vacuum was switched off as soon as bubbles exit the channel openings. This step was repeated 5 times. The remaining PVA was flushed out with a glucose solution, after which the GUV sample was pipetted in for sorting. Adapters were slowly inserted into the channel openings and then tubings (PTFE-tubing mm ID x 0.76 mm OD, Darwin Microfluidics) were connected to the adapters (Male Luer Fluid Connectors, Darwin Microfluidics) and the pump (G100-1L Peristaltic Pump for Incubator, Longer) sealed by two component glue (ecosil). The assembled chamber was left to equilibrate for 10 min. The sorting was then performed at a pumping speed of 0.2 rpm in up to 6 channels in parallel, which takes 3.3 min. The outgoing tubing is connected to a collection tube or another imaging channel.

## Acknowledgments

This work was funded by the European Union (ERC, ENSYNC, 101076997); the Deutsche Forschungsgemeinschaft (DFG, German Research Foundation) under Germany’s Excellence Strategy – EXC-3018/1 – 533587280, under TRR 392: Molecular evolution in prebiotic environments (Project number 521256690) and under SFB-1638/1–511488495 as well as the Human Frontiers Science Program (HFSP, reference number RGP003/2023). M.L. and K.G. thank the Hector Fellow Academy. We would like to acknowledge access to the infrastructure and support provided by the Cryo-EM Network at the Heidelberg University (HDcryoNet). We thank Claudia Helbig for technical support and the precision engineering workshop at Max Planck Institute for Medical Research in Heidelberg for manufacturing parts.

## Author contributions

M.L. developed, conducted and analysed most experiments and supervised M.U., who performed and analysed the centrifugation experiments and additional sedimentation experiments. S.M. developed the setup for DMD-based printing, T.A. implemented microwell printing. D.F. conducted electron microscopy imaging. M.L. and K.G. wrote the manuscript with input from all authors. K.G. supervised the research. Some items in the figures have been created with BioRender.com

## Additional information

Correspondence and requests for materials should be addressed to K.G.

## References

[1] Kriebisch, C. M. et al. A roadmap toward the synthesis of life. Chem 11 (2025).

[2] Abil, Z. & Danelon, C. Roadmap to building a cell: an evolutionary approach. Frontiers in Bioengineering and Biotechnology 8, 927 (2020).

[3] Van de Cauter, L., Van Buren, L., Koenderink, G. H. & Ganzinger, K. A. Exploring giant unilamellar vesicle production for artificial cells—current challenges and future directions. Small Methods 7, 2300416 (2023).

[4] Jalmar, O., García-Sáez, A. J., Berland, L., Gonzalvez, F. & Petit, P. Giant unilamellar vesicles (guvs) as a new tool for analysis of caspase-8/bid-fl complex binding to cardiolipin and its functional activity. Cell death & disease 1, e103–e103 (2010).

[5] Fujii, S., Matsuura, T., Sunami, T., Kazuta, Y. & Yomo, T. In vitro evolution of α-hemolysin using a liposome display. Proceedings of the National Academy of Sciences 110, 16796–16801 (2013).

[6] Tivony, R., Fletcher, M., Al Nahas, K. & Keyser, U. F. A microfluidic platform for sequential assembly and separation of synthetic cell models. ACS Synthetic Biology 10, 3105–3116 (2021).

[7] Bolognesi, G. et al. Sculpting and fusing biomimetic vesicle networks using optical tweezers. Nature communications 9, 1882 (2018).

[8] Wang, X., Tian, L. & Han, X. Response of giant unilamellar vesicles (guvs) in acoustic field. ChemSystemsChem 7, e202500014 (2025).

[9] Aparna, G. & Tetala, K. K. Recent progress in development and application of dna, protein, peptide, glycan, antibody, and aptamer microarrays. Biomolecules 13, 602 (2023).

[10] Sleath, H., Mognetti, B., Elani, Y. & Di Michele, L. Chemotactic crawling of multivalent vesicles along ligand-density gradients (2023).

[11] Giessler, F. et al. Growth, dissolution and segregation of genetically encoded rna droplets by ribozyme catalysis. Angewandte Chemie e19002 (2025).

[12] Sun, T., Kovac, J. & Voldman, J. Image-based single-cell sorting via dual-photopolymerized microwell arrays. Analytical chemistry 86, 977–981 (2014).

[13] Jia, H. et al. Shaping giant membrane vesicles in 3d-printed protein hydrogel cages. Small 16, 1906259 (2020).

[14] Ali, S. K. et al. Sedimentation-based confinement of individual giant unilamellar vesicles in microchamber arrays with a dynamically exchangeable outer medium. Advanced Materials Technologies 9, 2301976 (2024).

[15] Bartelt, S. M., Steinkühler, J., Dimova, R. & Wegner, S. V. Light-guided motility of a minimal synthetic cell. Nano Letters 18, 7268–7274 (2018).

[16] Dezi, M., Di Cicco, A., Bassereau, P. & Levy, D. Detergent-mediated incorporation of transmembrane proteins in giant unilamellar vesicles with controlled physiological contents. Proceedings of the National Academy of Sciences 110, 7276–7281 (2013).

[17] Tran, M. P. et al. Genetic encoding and expression of rna origami cytoskeletons in synthetic cells. Nature Nanotechnology 1–8 (2025).

[18] Jahnke, K. et al. Dna origami signaling units transduce chemical and mechanical signals in synthetic cells. Advanced Functional Materials 34, 2301176 (2024).

[19] Fragasso, A. et al. Reconstitution of ultrawide dna origami pores in liposomes for trans-membrane transport of macromolecules. ACS nano 15, 12768–12779 (2021).

[20] Thomsen, R. P. et al. A large size-selective dna nanopore with sensing applications. Nature communications 10, 5655 (2019).

[21] Krishnan, S. et al. Molecular transport through large-diameter dna nanopores. Nature communications 7, 12787 (2016).

[22] Burns, J. R. et al. Lipid-bilayer-spanning dna nanopores with a bifunctional porphyrin anchor. Angewandte Chemie (International ed. in English) 52, 12069 (2013).

[23] Dang, T. X., Hotze, E. M., Rouiller, I., Tweten, R. K. & Wilson-Kubalek, E. M. Prepore to pore transition of a cholesterol-dependent cytolysin visualized by electron microscopy. Journal of structural biology 150, 100–108 (2005).

[24] Jahnke, K., Grubmüller, H., Igaev, M. & Göpfrich, K. Choice of fluorophore affects dynamic dna nanostructures. Nucleic Acids Research 49, 4186–4195 (2021).

[25] Mohammed, A. M. & Schulman, R. Directing self-assembly of dna nanotubes using programmable seeds. Nano letters 13, 4006–4013 (2013).

[26] Burns, J. R., Seifert, A., Fertig, N. & Howorka, S. A biomimetic dna-based channel for the ligand-controlled transport of charged molecular cargo across a biological membrane. Nature nanotechnology 11, 152–156 (2016).

[27] Floroni, A. et al. Membraneless protocell confined by a heat flow. Nature Physics 1–8 (2025).

[28] Bell, N. A. & Keyser, U. F. Nanopores formed by dna origami: a review. FEBS letters 588, 3564–3570 (2014).

[29] Liu, F. et al. Engineering dna nanopores: from structural evolution to sensing and transport. Materials Today Bio 102137 (2025).

[30] Li, Y. et al. Leakless end-to-end transport of small molecules through micron-length dna nanochannels. Science Advances 8, eabq4834 (2022).

[31] Kuzuya, A., Wang, R., Sha, R. & Seeman, N. C. Six-helix and eight-helix dna nanotubes assembled from half-tubes. Nano letters 7, 1757–1763 (2007).

[32] Seifert, A. et al. Bilayer-spanning dna nanopores with voltage-switching between open and closed state. Acs Nano 9, 1117–1126 (2015).

[33] Göpfrich, K. et al. Large-conductance transmembrane porin made from dna origami. ACS nano 10, 8207–8214 (2016).

[34] Rausch, J. M. & Wimley, W. C. A high-throughput screen for identifying transmembrane pore-forming peptides. Analytical Biochemistry 293, 258–263 (2001).

[35] Zhang, J. et al. Microwell array chip-based single-cell analysis. Lab on a Chip 23, 1066–1079 (2023).

[36] Lai, R. L. & Huang, N.-T. Dimensional analysis and parametric studies of the microwell for particle trapping. Microfluidics and Nanofluidics 23, 121 (2019).

[37] Wu, H. et al. Facile method for fabricating microfluidic chip integrated with microwell arrays for cell trapping. Micromachines 10, 719 (2019).

[38] Yin, K. et al. Well-paired-seq: a size-exclusion and locally quasi-static hydrodynamic microwell chip for single-cell rna-seq. Small Methods 6, 2200341 (2022).

[39] Mohammed, A. M., Šulc, P., Zenk, J. & Schulman, R. Self-assembling dna nanotubes to connect molecular landmarks. Nature nanotechnology 12, 312–316 (2017).

[40] Ohi, M., Li, Y., Cheng, Y. & Walz, T. Negative staining and image classification—powerful tools in modern electron microscopy. Biological procedures online 6, 23–34 (2004).

[41] Punjani, A., Rubinstein, J. L., Fleet, D. J. & Brubaker, M. A. cryosparc: algorithms for rapid unsupervised cryo-em structure determination. Nature methods 14, 290–296 (2017).

[42] Van Heel, M., Harauz, G., Orlova, E. V., Schmidt, R. & Schatz, M. A new generation of the imagic image processing system. Journal of structural biology 116, 17–24 (1996).

[43] Liu, X. & Wang, H.-W. Single particle electron microscopy reconstruction of the exosome complex using the random conical tilt method. Journal of visualized experiments: JoVE 2574 (2011).

