## Supplementary Information for "Buoyancy-driven sorting of synthetic cells for nanopore activity"

#### Contents

**1 Supplementary Figures** **2**

**2 Supplementary Notes** **23**

**3 Supplementary Movies** **24**

### 1 Supplementary Figures

#### Supplementary Figure 1

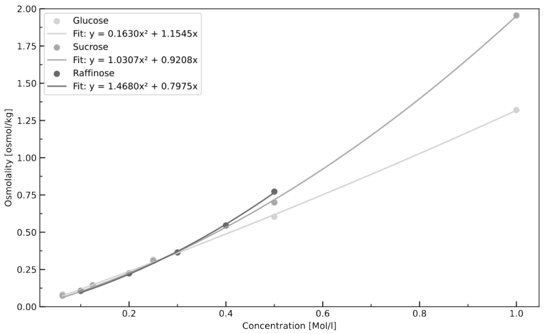


Figure S1: **Dependence of the osmolality on solute concentration for different sugars, fitted with a quadratic function ($y = ax^2 + bx$).** All sedimentation experiments were conducted at 280 mosmol/kg, corresponding to the osmolality of PBS (Osmomat 3000 basic, Gonotec GmbH). Deduced iso-osmotic concentrations are 235 mM glucose, 240 mM sucrose 243 mM raffinose.

#### Supplementary Figure 2

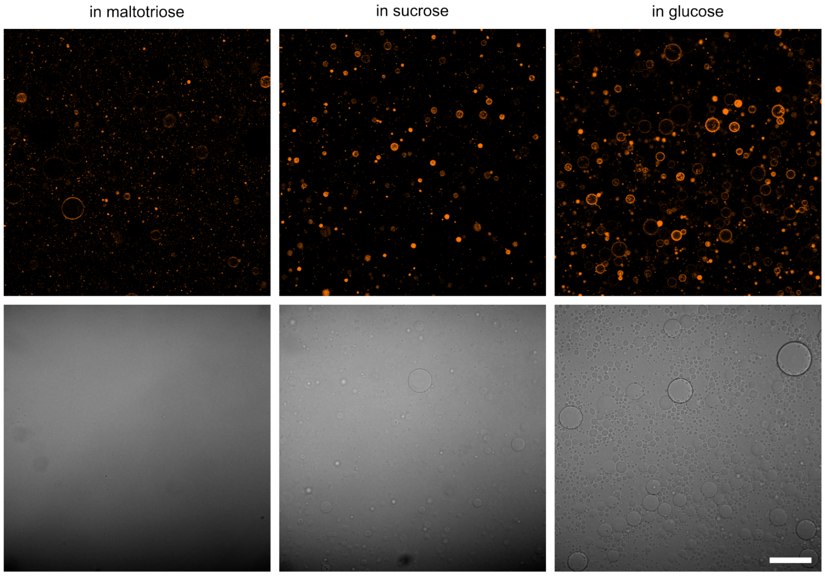


Figure S2: **Sedimentation of maltotriose-filled GUVs**. Confocal images of maltotriose-filled GUVs containing 1% Liss-Rhod PE (orange, $\lambda_{ex}$ = 561 nm) in different iso-osmotic sugar environments, as indicated (maltotriose $Mw_{malt}$ = 504 g/mol, sucrose $Mw_{sucr}$ = 342 g/mol and glucose $Mw_{gluc}$ = 180 g/mol). Fluorescence (top) and brightfield channels (bottom) are shown. Scale bar: 100 µm. GUVs sediment most in glucose, the lightest sugar environment.

#### Supplementary Figure 3

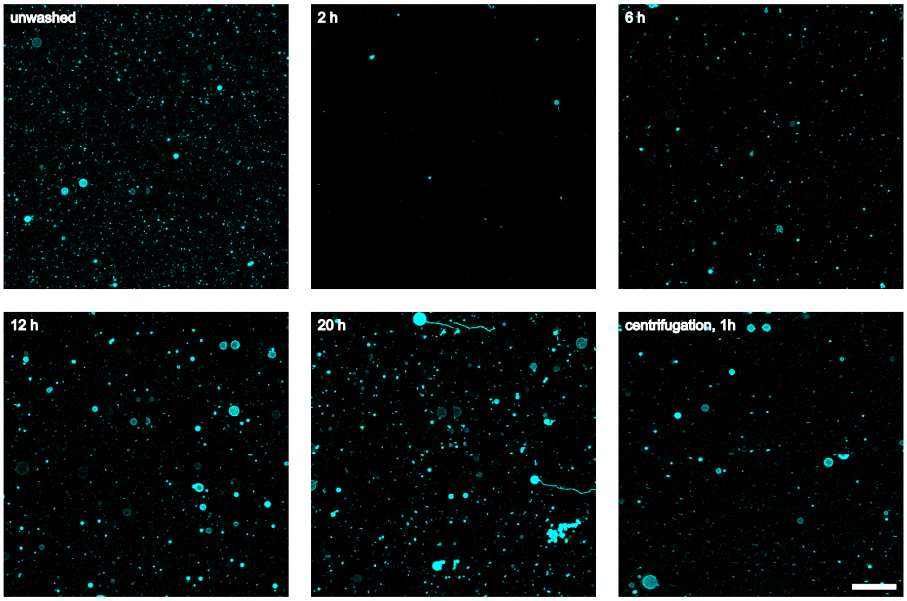


Figure S3: **Sedimentation as a washing protocol**. Confocal images of raffinose-filled GUVs (99 % DOPC, 1 % Atto488-DOPE, cyan, $\lambda_{ex} = 488$ nm) prior to the washing protocol (unwashed) and after sedimentation in iso-osmotic sucrose solutions for 2, 6, 12, and 20 h, as indicated. The number of GUV at the bottom of the observation chamber increases up to 12 h of sedimentation, followed by a pronounced decrease in GUV quality at longer times. Longer sedimentation times lead to co-sedimentation of small particles. Thus, 12 h of sedimentation is optimal to obtain a clean sample. Alternatively, centrifugation for 1 h at 100 g using a centrifuge with a rotor radius of 18.9 cm (Centrifuge 5810, S-4-104 Rotor, Eppendorf) can be applied to speed up the washing process. Scale bar: 100 µm.

#### Supplementary Figure 4

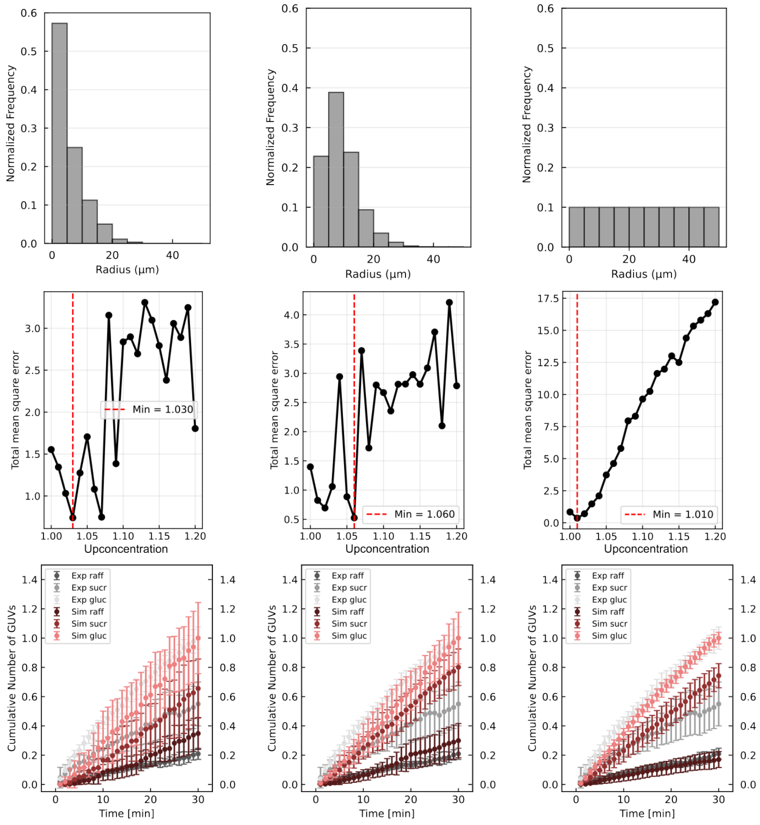


Figure S4: **Raffinose upconcentration during GUV formation. Upper row:** GUV radius distributions used for subsequent calculations as estimated from averaged radii of unwashed GUVs from the data shown in Fig. 2**e** (left), averaged radii of washed GUVs from the data shown in Figs. 3–5 (center), and a hypothetical (non-realistic) homogeneous GUV radius distribution used for comparison (right). **Middle row:** displays the deviation between the number of GUVs arrived at the bottom in simulation and experiment. It is calculated as the sum of the mean square errors, as a function of the assumed sugar upconcentration inside GUVs, for the respective GUV size distributions shown above. The upconcentration that fits best the experimental observation is indicated by a red dashed line. *[continues on next page]*

*[continued from previous page]*

**Bottom row:** normalised number of GUVs arriving at the bottom of the observation chamber, evaluated at upconcentration corresponding to the minimum total error (middle row). Experimental data is shown in shades of grey, simulation data is shown in shades of red for the different sugar environments. By comparing experiment and simulation data (according to Fig. 2**e** data, left column), we thus assume that the upconcentration is approximately 3 %, which lies in a range consistent with slightly varying GUV distributions (middle and right column).

#### Supplementary Figure 5

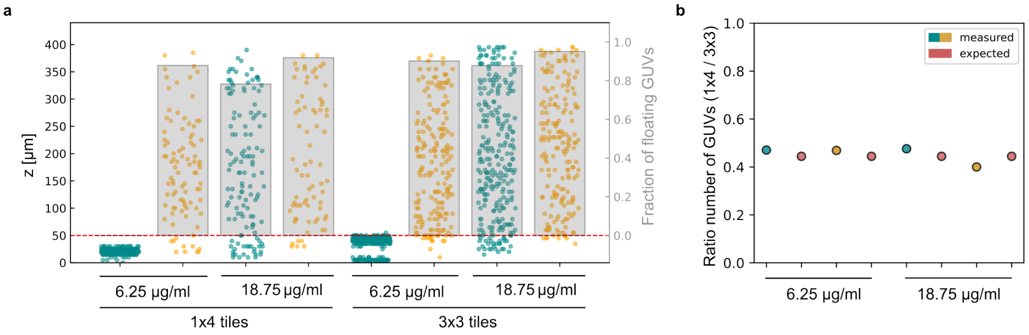


Figure S5: **The number of detected GUVs scales with the number of imaged tiles.** **a** Z-positions of GUVs without $\alpha$-hemolysin (cyan) and with 6.25 µg/ml monomers (27 nM pore) or 18.7 µg/ml monomers (81 nM pore) $\alpha$-hemolysin (orange) were measured over an imaging area comprising $1 \times 4$ tiles and compared to measurements of the same samples acquired over a larger $3 \times 3$ tile area. **b** Analysis of the ratio between measurements obtained from four and nine tiles shows that all values remain within 10% of the expected scaling, indicating consistent and tile-number–independent detection.

#### Supplementary Figure 6

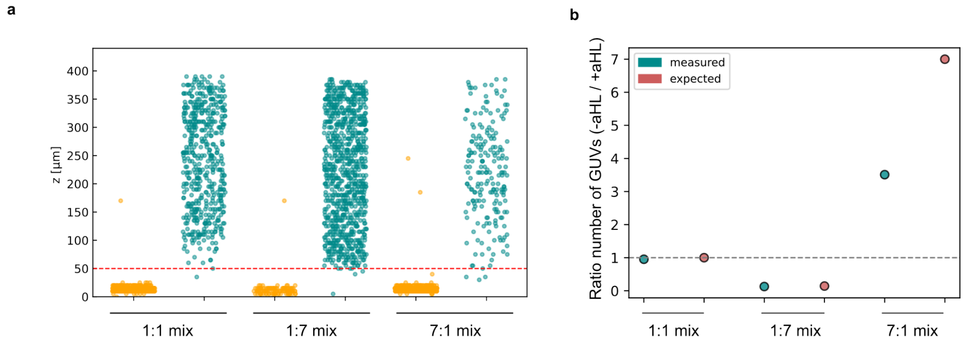


Figure S6: **Effect of different mixing ratios on sedimentation. a** The z-positions of mixed GUV populations without $\alpha$-hemolysin (orange) and with $\alpha$-hemolysin (cyan) were analysed at mixing ratios of 1:1, 1:7, and 7:1. **b** Analysis of the ratio between $-\alpha$-hemolysin and $+\alpha$-hemolysin measurements shows that the relative population fractions can be reliably quantified when GUV concentrations are medium to low or when GUVs are broadly distributed along the z-axis. Reduced precision is observed only at high fractions of $-\alpha$-hemolysin GUVs that accumulate in close proximity at the bottom of the imaging chamber, making the detection of GUVs more unreliable. This limitation does not affect the reliability of the sort, as population sorting does not require image analysis.

#### Supplementary Figure 7

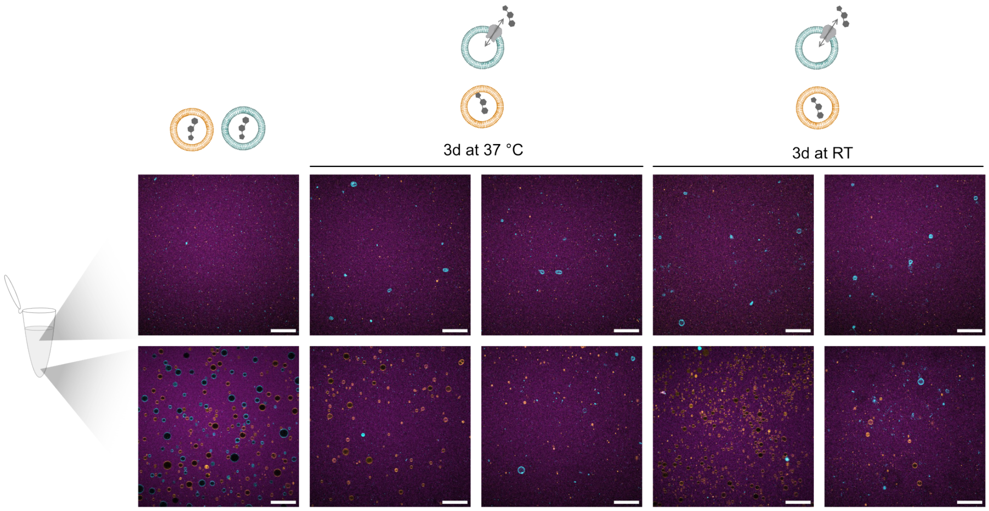


Figure S7: **Sedimentation assay in large volumes.** GUVs without pores settle to the bottom of the 1.5 ml reaction tube (first column, bottom). At the top of the tube no GUVs can be detected (first column, top). In case one population is incubated with pores (cyan, DOPC, 1% Atto488-DOPE, $\lambda_{ex} = 488\,\text{nm}$) after three days at 37 ° the permeabilised population can be clearly found in the upper part of the tube (second column, top), while the GUV population without pore (orange, DOPC, 1% Atto550-DOPE, $\lambda_{ex} = 561\,\text{nm}$) sinks to the bottom (second column, bottom). Also after three days at room temperature the same behaviour can be observed (third column). Scale bars: 100 µm. This shows that the sedimentation assay can, in principle, be scaled up for sorting of larger populations.

#### Supplementary Figure 8

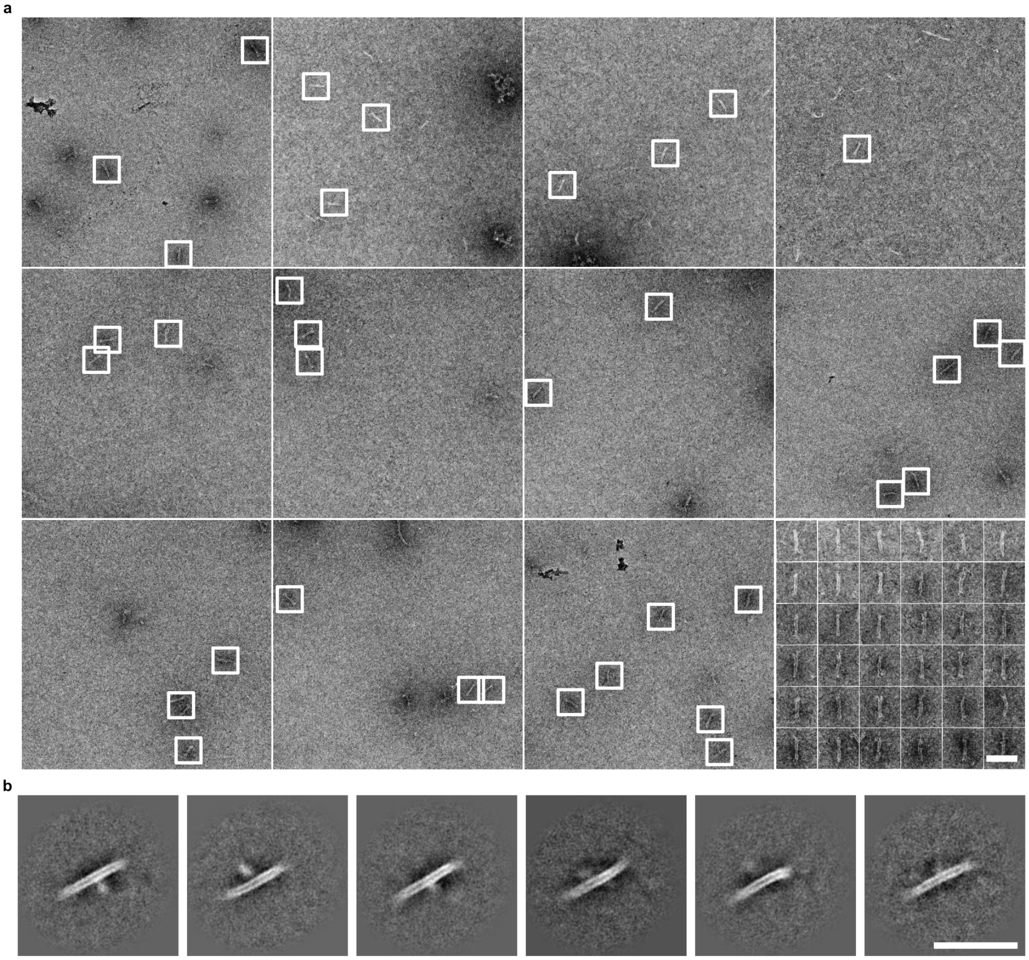


Figure S8: **Negative stain transmission electron microscopy images of the 12HB DNA origami pore.** **a** Overviews and selected particles (white frames) reprinted next to each other at the bottom right. Scale bar: 100 nm. **b** Class averages deduced from 600 overview micrographs and approximately 2500 particles. Scale bar: 100 nm.

#### Supplementary Figure 9

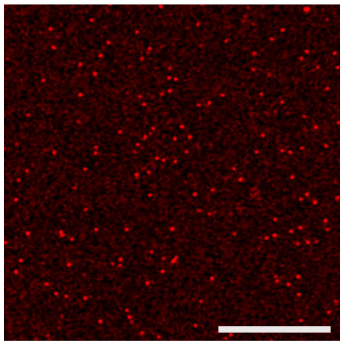

Figure S9: **12HB DNA origami pore characterization with superresolution microscopy.** Confocal airy scan micrograph of 12HB DNA origami pore labelled with 100 Atto647-modified staples positioned on the scaffold loop (red, $\lambda_{ex} = 640\,\text{nm}$) in bulk. Scale bar: 10 µm. In detail, on average 272 pores were counted per imaging area 77.34 µmx77.34 µm in a 90 µm high chamber in the Airy scan mode of the confocal microscope ($n = 18$ images), which translates to approx. 50 Mio. DNA origami pores per µl (0.47 ng/µL).

#### Supplementary Figure 10

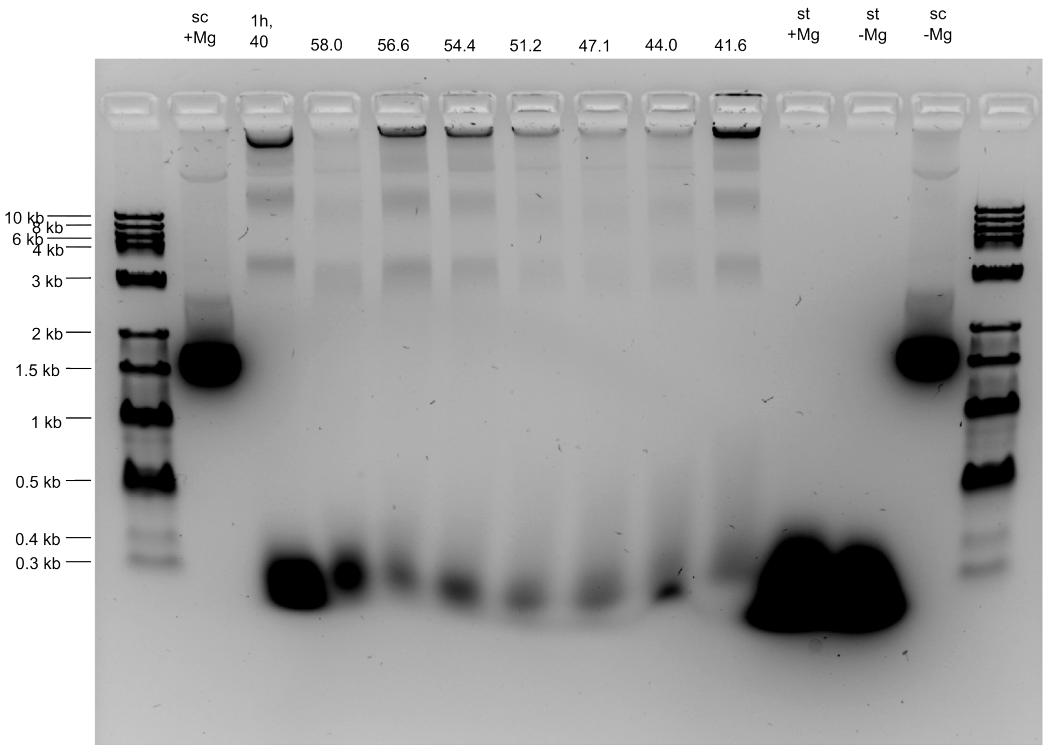


Figure S10: **Agarose gel electrophoresis comparing various 12HB DNA origami pore assembly conditions.** The lanes (left to right) contain: DNA ladder; 7249 bp scaffold in 5 mM $MgCl_2$; DNA origami assembled for 1 h at 40 °C; DNA origami assembled at 58 °C, 56.6 °C, 54.4 °C, 51.2 °C, 47.1 °C, 44 °C, and 41.6 °C; staple strands in 12 mM $MgCl_2$; staple strands without $MgCl_2$; 7249 bp scaffold without $MgCl_2$; and a second DNA ladder. Properly assembled 12HB DNA origami pores appear as a band above the scaffold.

#### Supplementary Figure 11

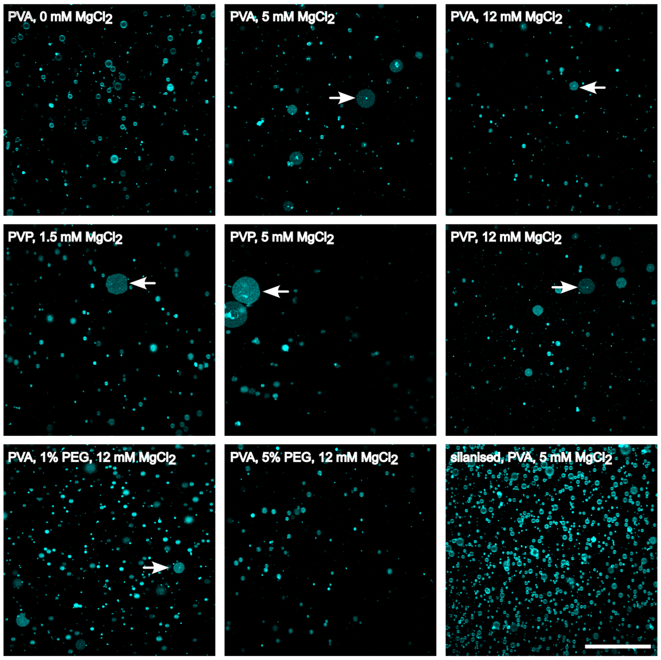


Figure S11: **Surface coating is critical when GUVs are denser than their surrounding medium.** In experiments using conventional PVA-coated slides without magnesium ions, GUVs behaved normally with no bursting observed (top left). However, at elevated magnesium concentrations, GUVs immediately bursted (white arrows) upon settling on the glass surface (top row). A similar behavior was observed on PVP-coated slides (middle row). When GUVs contained 1% PEGylated lipids, PVA coating still failed in the presence of 12 mM $MgCl_2$, whereas 5% PEG-ylated lipids provided partial protection against bursting (bottom row, left and center). For our standard lipid mixture (99 % DOPC, 1 % fluorescent DOPE), reliable prevention of bursting was achieved using silanised slides prepared for microwell fabrication, which allow covalent binding of the photoresist to the glass (bottom row, right). Scale bar: 200 µm.

#### Supplementary Figure 12

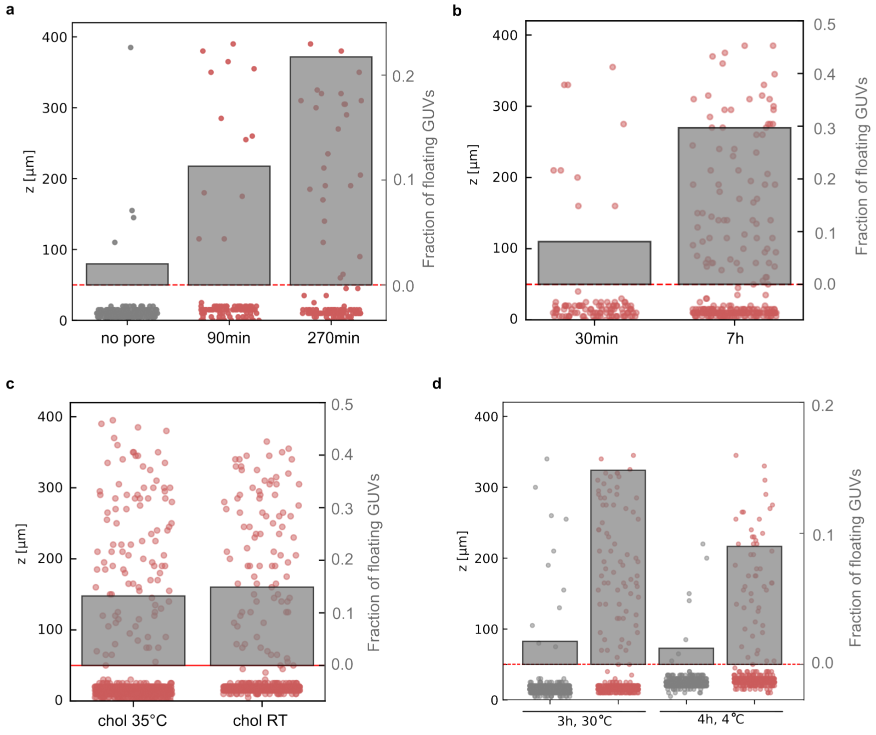


Figure S12: **12HB DNA origami pore insertion at longer incubation times and higher temperature.** Z-position of GUVs (left axis) and fraction of floating GUVs (right axis) without (gray) and with (red) 12HB DNA origami pores at different incubation times. **a** GUVs were monitored without and with 7nM 12HB DNA origami pore after 90 min and 270 min incubation. **b** GUVs were monitored with 22 nM 12HB DNA origami pore after 30 min and 7 h incubation. **c** 12HB DNA origami pores were incubated with cholesterol-tagged DNA at 35 °C and at room temperature prior to addition to the GUVs and then monitored after 2h incubation at room temperature. **d** GUVs were incubated for 3 h at 30 °C and for 4 h at 4 °C. Overall, higher DNA origami pore concentrations, longer incubation times and elevated temperatures yield a higher fraction of floating GUVs.

#### Supplementary Figure 13

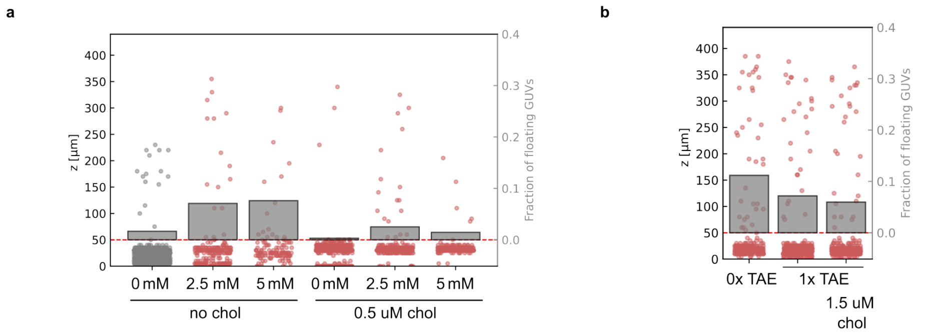


Figure S13: **Control experiments with magnesium ions and cholesterol-tagged DNA. a** GUVs in the absence of magnesium ions and pores do not exhibit floating. At 2.5 mM or 5 mM $MgCl_2$, incubation at room temperature for 1 h results in a measurable fraction of floating GUVs. This effect is reduced in the presence of cholesterol-tagged DNA. **b** The presence of 5 mM $MgCl_2$, incubation at room temperature for 2 h results in a measurable fraction of floating GUVs. Addition of 1× TAE buffer attenuates this effect. It is further reduced in the presence of cholesterol-tagged DNA.

### Supplementary Figure 14

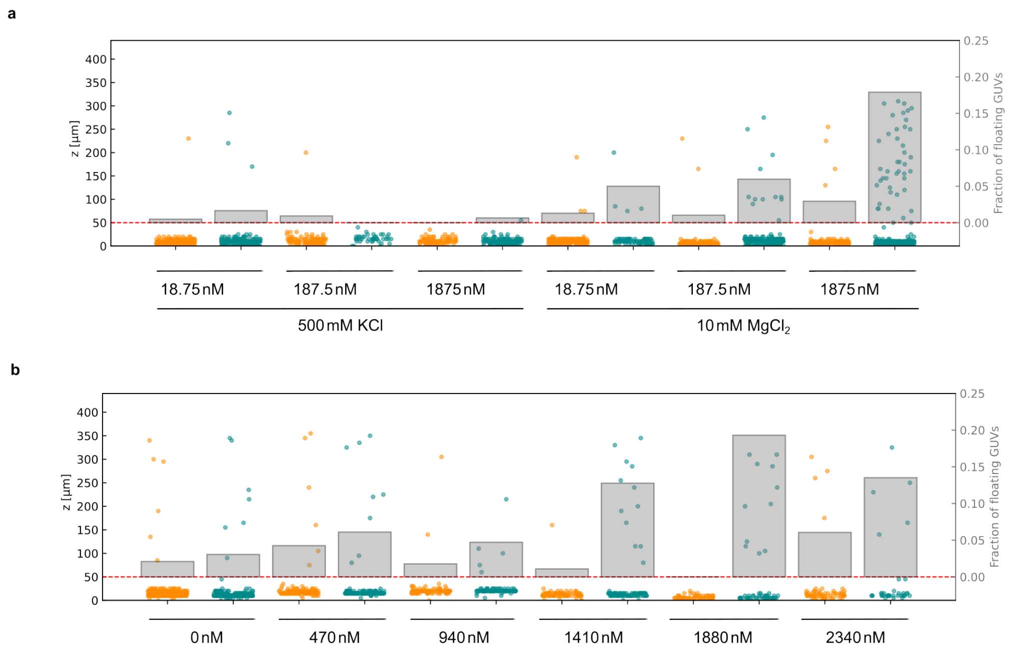


Figure S14: **Sedimentation assay with 6-helix bundle (6HB) DNA tile pore.** Z-position of GUVs without (orange) and with 6HB DNA tile pore (cyan) and fraction of floating GUVs. **a** Effect of pore concentration in 500 mM KCl and 10 mM $MgCl_2$. **b** Effect of pore concentration at 10 mM $MgCl_2$.

#### Supplementary Figure 15

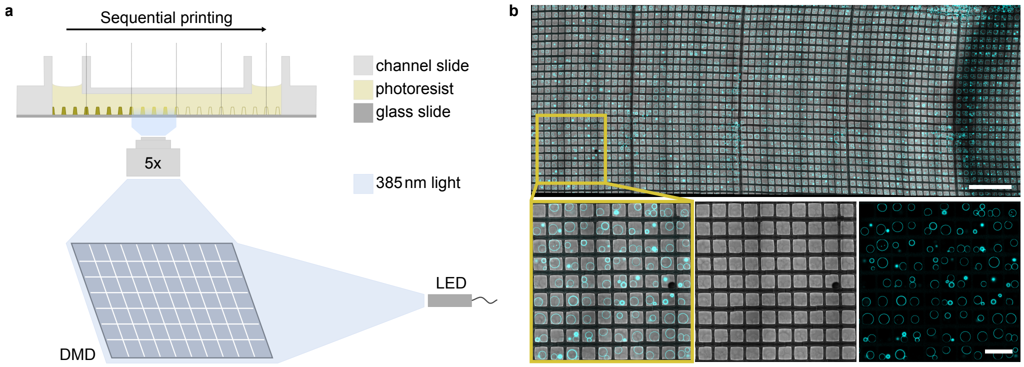


Figure S15: **Microwells inside of a multichannel chamber for trapping GUVs during sorting.** **a** Schematic illustration of microwell fabrication by a 385 nm LED coupled into a digital micromirror device, which reflects the light according to a digital mask. A 5x objective focuses the reflected light into a liquid photoresist inside of a channel of a multichannel slide, resulting in spatially confined polymerisation of microwell walls. After sequential printing in a tile-by-tile manner, uncured photoresist is being washed out. **b** Composite image of GUVs (cyan, 99 % DOPC, 1 % Atto488-DOPE, $\lambda_{ex} = 488$ nm) settled due to a density gradient and trapped inside of printed microwells (gray) inside of a multichannel slide. Zoom image shows GUVs inside of the microwells as overlay and individual channels (bottom). Scale bars: 1 mm (top) and 200 µm (bottom).

#### Supplementary Figure 16

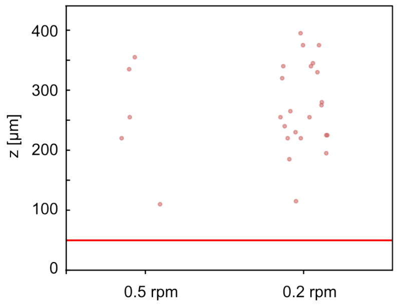


Figure S16: **Effect of pumping speed on GUV harvesting.** A more than 4-fold increase in harvested GUVs was achieved for slower pumping speeds (0.2 rpm, 3.3 min for 40 µl) compared to higher ones (0.5 rpm, 1.5 min for 40 µl), presumably due to GUV disruption at higher speeds.

#### Supplementary Figure 17

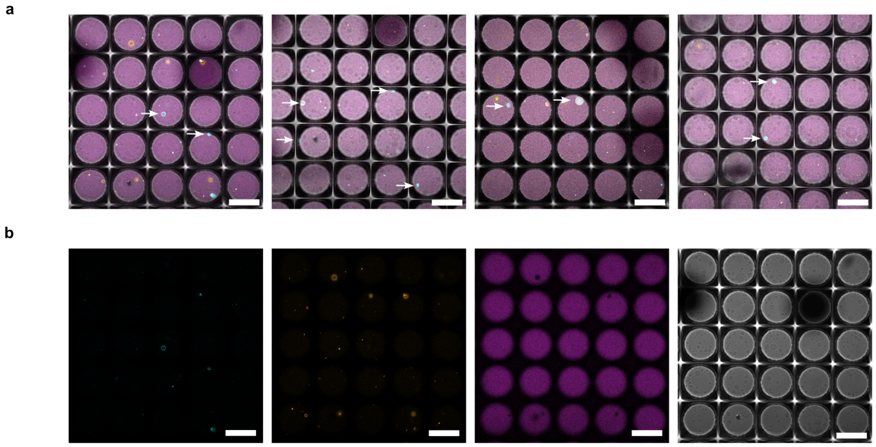


Figure S17: **Localisation of GUVs with and without $\alpha$-hemolysin pore in the microwells. a** Confocal overlay images of four channels (GUVs without $\alpha$-hemolysin: orange, $\lambda_{ex} = 561\,nm$; GUVs with $\alpha$-hemolysin: cyan, $\lambda_{ex} = 488\,nm$; microwell structure: grey, brightfield; Alexa Fluor 647 influx dye: purple, $\lambda_{ex} = 640\,nm$). The four images show four different positions. **b** Split channel view of the first image from above. Scale bars: 100 µm.

#### Supplementary Figure 18

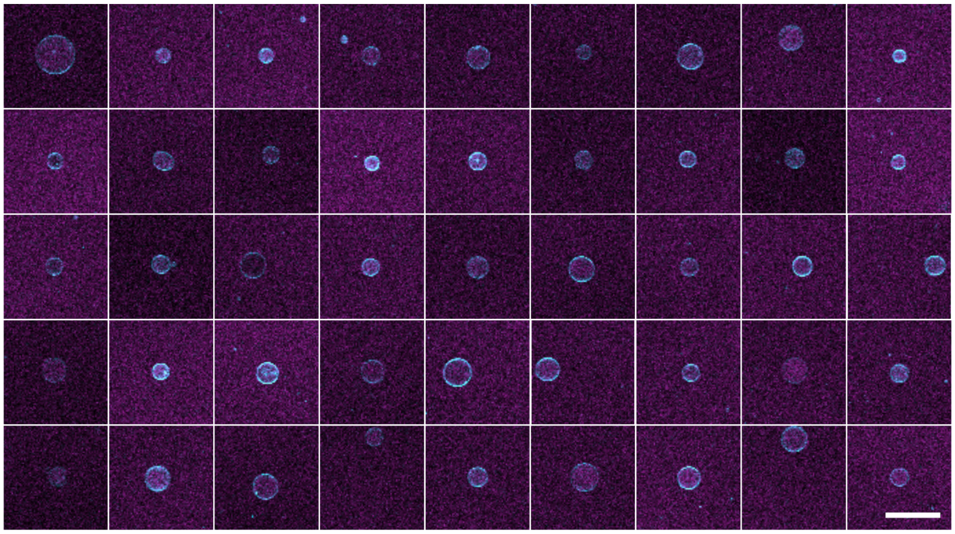

Figure S18: **Harvested GUVs are intact after sorting.** Confocal overlay images of sorted GUVs (cyan, 99 % DOPC, 1 % Atto488-DOPE, $\lambda_{ex}$ = 488 nm) immersed in a solution containing Alexa Fluor 647 to visualise dye influx (purple, $\lambda_{ex}$ = 640 nm). Scale bar: 100 µm. 98 % of harvested GUVs show dye influx.

#### Supplementary Figure 19

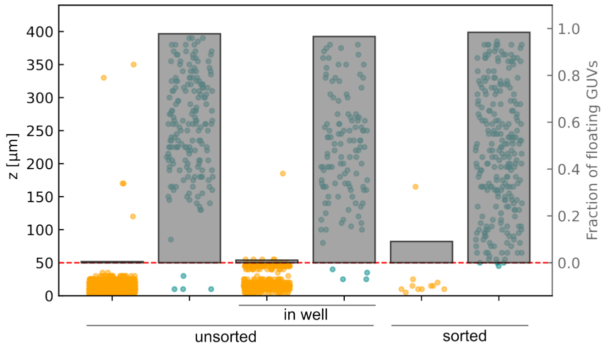


Figure S19: **GUVs containing $\alpha$-hemolysin (cyan, 99 % DOPC, 1 % Atto488-DOPE, $\lambda_{ex} = 488\,\mathrm{nm}$) can be distinguished and separated from pore-free GUVs (orange, 99 % DOPC, 1 % Atto550-DOPE, $\lambda_{ex} = 561\,\mathrm{nm}$).** The z-position and the fraction of floating GUVs are shown for the unsorted population in a plain imaging chamber, in microwell structures, and for the sorted population.

#### Supplementary Figure 20

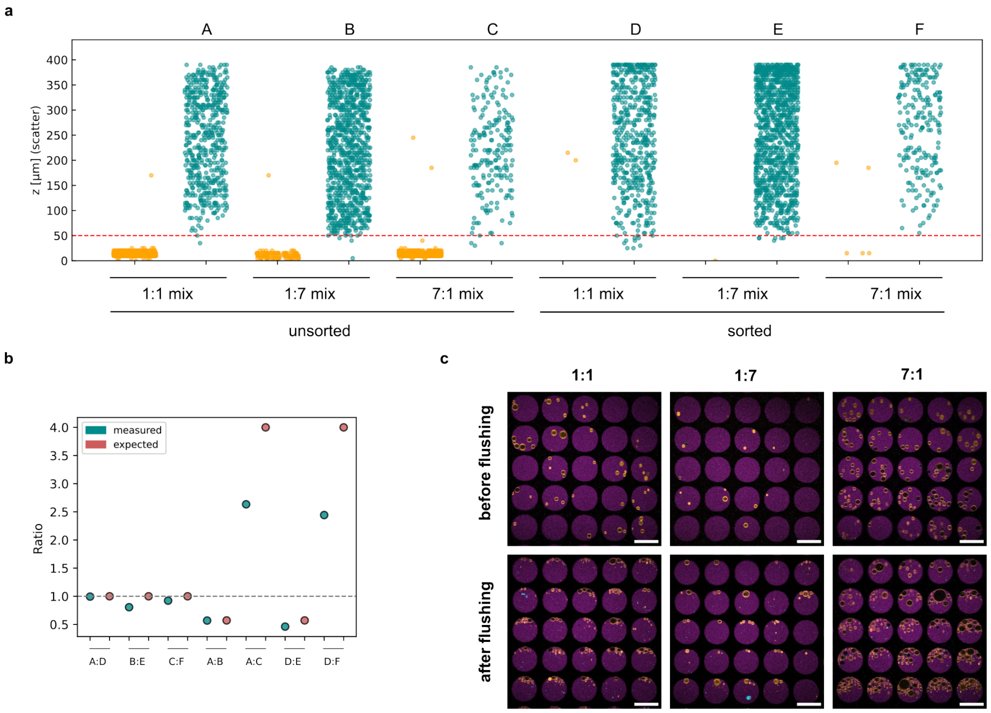


Figure S20: **Harvesting different fractions of pore-active GUVs.** Different mix fractions (1:1, 1:7 and 7:1) of GUV populations without pore (orange, 99 % DOPC, 1 % Atto550-DOPE, $\lambda_{ex} = 561$ nm) and with pore (cyan, 99 % DOPC, 1 % Atto488-DOPE, $\lambda_{ex} = 488$ nm) are prepared and measured before and after sorting. **a** The z-position and the fraction of floating GUVs are shown for the unsorted populations as well as the sorted populations. **b** Measured and expected pore-containing GUV count ratios of different conditions (mix ratios, before and after sorting). The sorting assay is capable of sorting a small population of 12.5 % (7:1 mix) and a large population of 87.5 % (1:7 mix) maintaining the integrity of the GUVs after sorting. The high deviation from the expected ratio in A:C presumably results from an unprecise mixture due to varying yields of the two GUV batches. It is important to note that the same can be observed for D:F confirming the high performance of the sorting assay. **c** Confocal overlay images of sedimented GUV populations at the bottom of the microwells of different mixes before and after flushing. Scale bars: 100 µm.

### 2 Supplementary Notes

#### Supplementary Note 1: Assumptions for simulation of GUV sedimentation

To calculate the accumulation of GUVs at the bottom, we assume the GUV as a sphere of mass $m = \frac{4}{3}\pi r^3 \rho_{GUV}$ and buoyancy $F_b$ acts against gravity $F_g$. So the downward acting force is given by the buoyancy-corrected gravity $g'$ as $F_{down} = mg' = mg(1 - \frac{\rho_{med}}{\rho_{GUV}})$. The Reynolds number calculates as $Re \approx \frac{2rv_e\rho_{med}}{\eta}$. For $r = 50\,\mu\text{m}$, $v = 1\,\text{cm/h}$, $\rho_{med} \approx 1030\,\text{kg/m}^3$, $\eta_{med} \approx 1.2\,\text{Pas}$ it reveals $Re << 1$, a linear Stokes regime as usual for water. In a classical Stokes regime, we assume $F_{fric} = 6\pi\eta r v = Kv$ and for $F_{fric} = F_{down}$, we get $Kv = mg'$ , so the end velocity of the GUV is deduced $v_e = \frac{mg'}{K} = \tau g'$. The z-position over time reads as $z(t) = z_0 - (v_e t) + v_e\tau(1 - e^{-t/\tau})$ with $\tau = m/6\pi\eta r$, $\tau < 10^-6$ so $t >> \tau$ and the exponential term is irrelevent for observed time scales.

The calculations take into account different randomly distributed starting z-positions (in case of channel sedimentation $z_{\text{max}}$ =400 µm), and a GUV radii distribution of up to 50 µm as experimentally observed. The effect of different GUV radii distributions is shown in Supplementary Fig. 4. The densities of 280 mosmol/kg sugar solutions are measure as $\rho_{\text{raff}}$ =1058 kg/m$^3$, $\rho_{\text{sucr}}$ =1032 kg/m$^3$, $\rho_{\text{gluc}}$ =1023 kg/m$^3$ and viscosities of the same are deduced from literature as $\eta_{\text{raff}} = 1.455$ mPas, $\eta_{\text{sucr}} = 1.156$ mPas, $\eta_{\text{gluc}} = 1.073$ mPas, [1] [2]. An additional lipid density of 0.696 kg/m$^3$ is applied. The simulation is calculated for 30 min, one time point per minute. The simulation methods are provided in the Methods section.

### 3 Supplementary Movies

#### Supplementary Movie 1: 3D-reconstructed time series of GUV sedimentation

Two populations of GUVs without $\alpha$-hemolysin (orange, 99 % DOPC, 1 % Atto550-DOPE $\lambda_{ex}$ = 561 nm) and with $\alpha$-hemolysin (cyan, 99 % DOPC, 1 % Atto488-DOPE, $\lambda_{ex}$ = 488 nm) are mixed in a 1:1 ratio and imaged in a 400 µm high channel. After storage upside-down, the sedimentation behaviour was imaged with a LSM 910 Lightfield 4D (Zeiss). A composite 3D reconstruction time series is shown. Box size: 1400 µm x 1400 µm x 400 µm.

#### Supplementary Movie 2: 3D-reconstructed time series of GUV sedimentation, pore-active population

Separate channel display of Supplementary Movie 1, showing only one channel with $\alpha$-hemolysin (cyan, 99 % DOPC, 1 % Atto488-DOPE, $\lambda_{ex}$ = 488 nm).

#### Supplementary Movie 3: 3D-reconstructed time series of GUV sedimentation, pore-inactive population

Separate channel display of Supplementary Movie 1, showing only one channel without $\alpha$-hemolysin (orange, 99 % DOPC, 1 % Atto550-DOPE, $\lambda_{ex}$ = 561 nm).

#### Supplementary Movie 4: Time series of 3D-lumen of artificial populations, Technical repeat

Another mix has been prepared and imaged as in Supplementary Movie 1.

#### Supplementary Movie 5: Time series of dye influx and GUV floating with 12HB DNA origami pores

Combined time series of four different positions of GUVs (cyan, 99 % DOPC, 1 % Atto488-DOPE, $\lambda_{ex}$ = 488 nm) with DNA origami pores and immersed in a solution containing Alexa Fluor 647 to probe dye influx (purple, $\lambda_{ex}$ =

640 nm). The Movie shows dye influx followed by floating of some of the GUVs over time (indicative of pore insertion). Scale bar: 100 µm.

##### Supplementary Movie 6: 3D reconstruction of the microwell architecture with GUVs

Rotation angle 90° of a 3D reconstruction of microwells (gray) filled with GUVs (orange, 99 % DOPC, 1 % Atto550-DOPE, $\lambda_{\mathrm{ex}} = 561$ nm). Scale bar: 100 µm.

##### Supplementary Movie 7: Layering of GUVs in microwell

Z-stack of one exemplary microwell filled with several GUVs immersed in a solution containing Alexa Fluor 647 to probe dye influx (purple, $\lambda_{ex} = 640$ nm). GUVs of one population (orange, 99 % DOPC, 1 % Atto550-DOPE, $\lambda_{ex} = 561$ nm, unfilled) appear at the bottom, others (cyan, 99 % DOPC, 1 % Atto488-DOPE, $\lambda_{ex} = 488$ nm, filled) in upper layers. Although the permeated GUVs are only positioned at the top of the well, a slight penetration of the applied flow into the well is sufficient to extract them and carry them downstream while the flow is not strong enough to perturb GUVs at the bottom. Scale bar: 100 µm.

#### References

[1] Chirife, J. & Buera, M. A simple model for predicting the viscosity of sugar and oligosaccharide solutions. *Journal of Food Engineering* **33**, 221–226 (1997).

[2] Washburn, E. W. & Williams, G. Y. The viscosities and conductivities of aqueous solutions of raffinose. *Journal of the American Chemical Society* **35**, 750–754 (1913).
